# Uncovering Convergent Cell State Dynamics Across Divergent Genetic Perturbations Through Single-Cell High-Content CRISPR Screening

**DOI:** 10.1101/2025.05.02.651939

**Authors:** Zihan Xu, Ziyu Lu, Aileen Ugurbil, Abdulraouf Abdulraouf, Andrew Liao, Jianxiang Zhang, Wei Zhou, Junyue Cao

**Author notes:** Correspondence (Z.X.), (J.C.).

## Abstract

High-throughput genomic studies have uncovered associations between diverse genetic alterations and disease phenotypes; however, elucidating how perturbations in functionally disparate genes give rise to convergent cellular states remains challenging. Here, we present ***PerturbFate***, a high-throughput, cost-effective, combinatorial-indexing single-cell platform that enables systematic interrogation of massively parallel CRISPR perturbations across the full spectrum of gene regulation, from chromatin remodeling and nascent transcription to steady-state transcriptomic phenotypes. Using *PerturbFate*, we profiled over 300,000 cultured melanoma cells to characterize multi-modal phenotypic and gene regulatory responses to perturbations in more than 140 Vemurafenib resistance-associated genes. We uncovered a shared dedifferentiated cell state marked by convergent transcription factor (TF) activity signatures across diverse genetic perturbations. Combined inhibition of cooperative TF hubs effectively reversed cellular adaptation to Vemurafenib treatment. We further dissected phenotypic responses to perturbations in Mediator Complex components, linking module-specific biochemical properties to convergent gene activations. Together, we reveal common regulatory nodes that drive similar phenotypic outcomes across distinct genetic perturbations. We also delineate how perturbations in functionally unrelated genes reshape cell state. *PerturbFate* thus establishes a versatile platform for identifying key molecular regulators by anchoring multi-modal regulatory dynamics to disease-relevant phenotypes.

## *PerturbFate* for capturing multi-layer single-cell readout upon genetic perturbations

High-throughput genomic sequencing has revolutionized our ability to identify genetic regulators of disease phenotypes, from oncogenesis(*1*), drug adaptation(*2*), to neurodegeneration(*3*). For example, unbiased bulk CRISPR screens have nominated genes whose loss- or gain-of-function confers resistance to targeted therapies, yet these hits span molecular processes as diverse as epigenetic modification, transcriptional regulation, and signaling cascades(*4*). Such diversity raises a fundamental question: how do these biologically diverse perturbations result in the similar phenotypic outcome? Do these disparate upstream events converge on a common downstream genetic or epigenetic program that ultimately induce the phenotypic change? Uncovering this shared program is essential for better understanding disease associated cellular states across a broad spectrum of genetic backgrounds.

Single-cell CRISPR screening approaches combining pooled perturbations with rich phenotypic readouts offer an unique opportunity into convergent molecular states across diverse genetic perturbations(*5–8*). However, most existing methods capture only a static snapshot of single molecular layer, preventing a comprehensive view of genetic perturbations-induced gene regulatory responses. In addition, reliance on droplet-based platforms limits both throughput and cost-efficiency(*9*), constraining our ability to map shared regulatory programs across large perturbation libraries.

To overcome these challenges, we developed *PerturbFate*, a highly scalable single-cell combinatorial indexing platform with flexible multi-modal readout that enables concurrently profiling chromatin accessibility, nascent and pre-existing transcriptomes, or steady-state transcriptomes, together with sgRNA identity, in individual cells. In brief (**Fig. 1A**), the workflow comprises: 1) Pooled sgRNA transduction: Cells are transduced with a pooled sgRNA Lentivirus library, generating a heterogeneous cell population in which every subpopulation harbors individual gene knockdowns. 2) Nascent transcript labeling: before harvesting cells, 5-Ethynyl Uridine (5-EU) is added to the culture medium to mark newly synthesized transcripts(*10*). 3) Chromatin tagmentation: After harvesting, fixation, and permeabilization, Tn5-mediated transposition assay for transposase-accessible chromatin (ATAC) is carried out in bulk, tagging open chromatin regions. 4) Reverse transcription is performed in bulk to capture both newly synthesized and pre-existing RNA transcripts, including sgRNA sequences. 5) Indexing ligation: Cells undergo two rounds of split-and-pool steps for indexing ligation reactions, introducing cell barcode combinations to RNA and ATAC fragments. 6) Click chemistry is performed in bulk on permeabilized post-ligation cells to conjugate biotin to 5-EU-labeled cDNA–RNA hybrids. 7) Streptavidin pull-down: following cell lysis, biotin-conjugated nascent cDNA–RNA hybrids with the first two cell barcodes are isolated using streptavidin beads. 8) Library preparation: After second-strand synthesis of cDNA and Read2 tagmentation, multiplex indexing PCR and sgRNA enrichment PCR are performed sequentially to amplify and separate multi-omics sequencing libraries. 9) Single-cell multi-modal assignment: Throughout this workflow, each modality in each single cell is linked to a unique combination of DNA barcodes. Computational assignment of sequencing reads and matching of multiple modalities are performed to recover single cells *in silico* (**Fig. S1**).

**Fig. 1.**
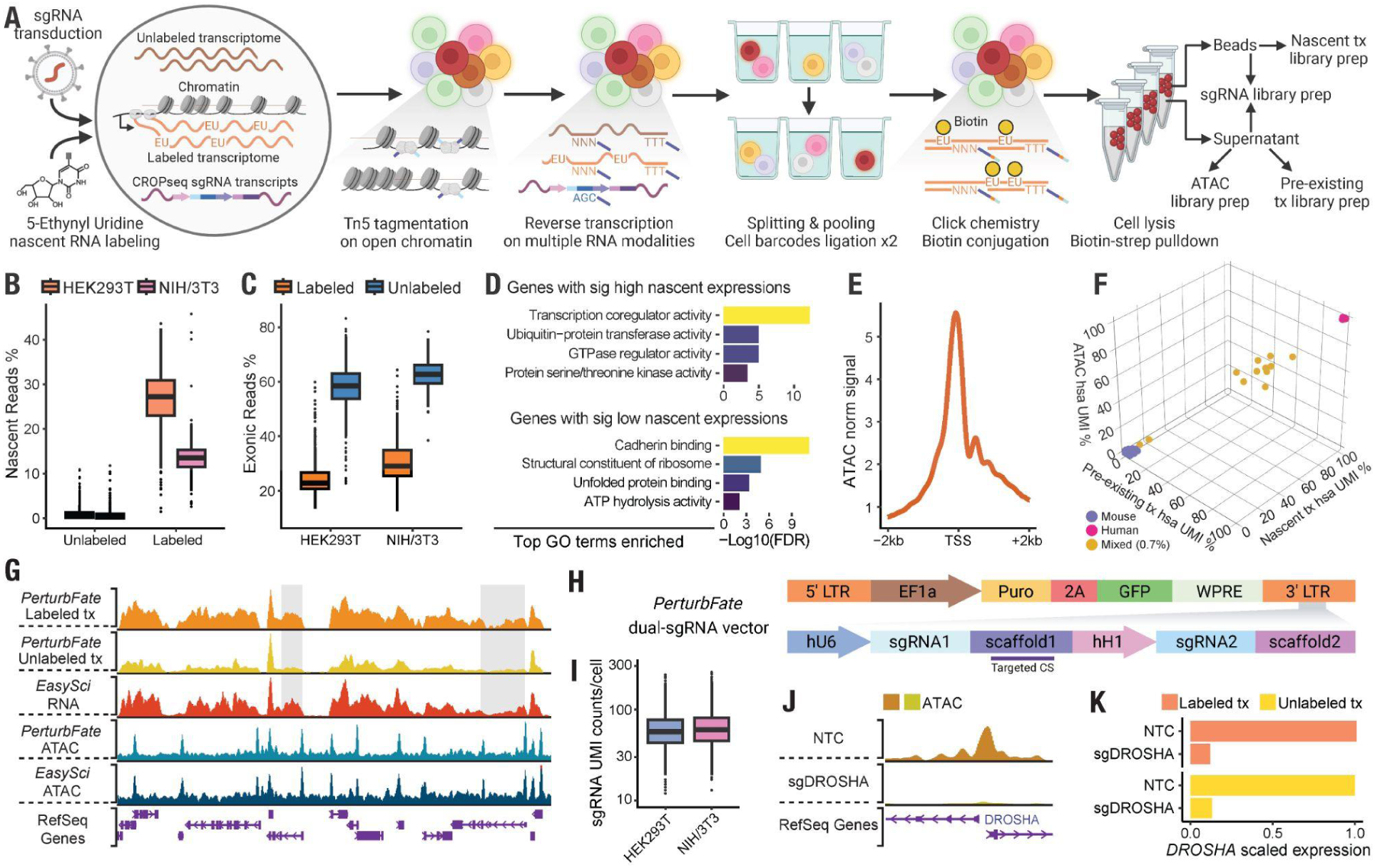
*PerturbFate* enables simultaneous capture of nascent and pre-existing transcriptomes, chromatin accessibility, and sgRNA identities from the same single cell. **(A)** Schematic of the *PerturbFate* library preparation workflow for co-capturing nascent and pre-existing transcriptomes, chromatin accessibility, and sgRNA identities in individual cells. **(B)** Box plot showing the percentage of labeled nascent gene count UMIs detected in 5-EU-labeled and unlabeled cells (HEK293T unlabeled *n* = 867, HEK293T labeled *n* = 651, NIH/3T3 unlabeled *n* = 289, NIH/3T3 labeled *n* = 206). **(C)** Box plot showing the exonic proportion of reads from single-cell labeled nascent RNA and unlabeled pre-existing RNA (HEK293T *n* = 1,246, NIH/3T3 *n* = 395). **(D)** Bar plots showing the top enriched Gene Ontology (GO) terms in genes with significantly higher (upper panel) or lower (lower panel) nascent counts, as identified by single-cell differential expression analysis comparing nascent and pre-existing transcriptomes. **(E)** Line plot showing the normalized ATAC read coverage enrichment around TSS. For HEK293T cells profiled by *PerturbFate*, ATAC reads were aggregated in 100 bp bins within a 4 kb window and normalized by the mean read coverage in the first and last 100 bp bins. **(F)** 3D dot plot showing the consistency of species assignment across single-cell molecular modalities. Human HEK293T cells and mouse NIH/3T3 cells were mixed for *PerturbFate* profiling, and sequencing reads from each layer were aligned to a human- mouse hybrid reference genome to examine the purity of single-cell data. **(G)** Genome tracks displaying signal densities of different molecular modalities in HEK293T cells profiled by *PerturbFate* or by single-modality methods in a representative genomic region. Signals were normalized by counts per million (CPM) and visualized using the WashU Epigenome Browser(*17*). **(H)** Schematic of the optimized *PerturbFate* dual-sgRNA expression construct. An mRNA encoding anti-Puromycin and EGFP tags, along with sgRNA sequences at the 3′ end, is driven by the Pol II promoter; a pair of sgRNAs is separately driven by the hU6 and hH1 promoters. **(I)** Box plot showing single-cell sgRNA UMIs obtained from human (HEK293T, *n* = 1,219) and mouse (NIH/3T3, *n* = 827) cells. **(J)** Genome tracks showing aggregated ATAC chromatin accessibility around the *DROSHA* TSS in NTC (non-targeting control) cells and in *DROSHA*-knockdown cells profiled by *PerturbFate* (normalized by CPM). **(K)** Bar plots showing normalized pseudobulk expression levels of *DROSHA* in NTC and *DROSHA*-knockdown cells. Single-cell nascent and pre-existing transcriptome counts were combined by perturbation and normalized by CPM; the expression levels of *DROSHA* in each layer were scaled to the expression level in NTC cells. Boxes in box plots indicate the median and interquartile range (IQR), with whiskers indicating 1.5× IQR.

We extensively optimized key steps (*e.g.,* fixation, tagmentation, RT) to enhance the efficiency and robustness of *PerturbFate* (**Fig. S1-2**). For example, by lowering the temperature for reverse transcription after ATAC, the quality of chromatin accessibility modality was substantially improved (**Fig. S2A**); by testing different fixation conditions and reverse transcription recipes, we extensively enhanced the cell recovery and RNA capture sensitivity (**Fig. S2B-F**). Specifically, to minimize sgRNA off-target effect and heterogeneous perturbation responses caused by different sgRNAs targeting the same gene, we further engineered the *CROPseq-OPTI* vector used in our previous study(*8*, *11*, *12*) by incorporating two sgRNA expression cassettes, driven by the hU6 and hH1 promoters, into the 3′ long terminal repeat (LTR) of the lentiviral construct (**Fig. S1A**)(*13*). We also introduced a variant sgRNA scaffold containing a capture sequence downstream of sgRNA-1(*14*), enabling the direct sequencing of the sgRNA-1 sequence for target gene identification (**Fig. S1A**). Moreover, *PerturbFate* is fully compatible with parallel bulk CRISPR screening and we adapted the workflow to co-capture single-cell steady-state transcriptome and chromatin accessibility in response to pooled CRISPR perturbations (**Fig. S3**).

We first validated the ability of *PerturbFate* to recover nascent transcriptomics at single-cell resolution by profiling a mixture of human HEK293T and mouse NIH/3T3 cells, with or without a one-hour 5-EU labeling. As expected, we detected minimal background nascent reads in unlabeled controls with a median of 0.38% in HEK293T and 0.29% in NIH/3T3, compared to the 27.2% and 13.5% of nascent read counts recovered from labeled HEK293T and NIH/3T3 cells, respectively (**Fig. 1B**). Also, a significantly lower proportion of reads from labeled transcriptomes aligned to gene exons compared to unlabeled controls (**Fig. 1C**). Consistent with previous studies(*8*, *15*), genes with high nascent read counts were enriched in dynamic biological processes, including transcriptional coregulation and GTPase regulator activities; genes with slower RNA turnover (“low nascent reads”) were strongly associated with housekeeping functions such as cadherin binding and ribosomal processes (**Fig. 1D**).

Moreover, the strong enrichment of ATAC signals centered at transcription start sites (TSS) proved the high efficiency of capturing chromatin accessibility by *PerturbFate* (**Fig. 1E**). The species-mixing experiment revealed highly consistent single-cell purity across modalities, with only 0.7% of cells displaying undetermined species origin (**Fig. 1F**). In addition, the chromatin accessibility and transcriptome profiles obtained by *PerturbFate* showed strong concordance with those generated using single-modality methods (**Fig. 1G**). Notably, the labeled nascent RNA layer in *PerturbFate* showed a higher coverage toward the 5′ ends and intronic regions of genes compared with the non-labeled transcriptome, supporting its unique capacity to capture newly synthesized, non-spliced transcripts (**Fig. 1G**).

We next evaluated the sgRNA capture efficiency of *PerturbFate*. HEK293T and NIH/3T3 cells were transduced with species-specific sgRNAs at a low multiplicity of infection (MOI = 0.2) via the optimized dual-sgRNA system (**Fig. 1H**), followed by *PerturbFate* profiling of the cell mixture. We obtained 98.1% of cells with sgRNAs detected, in which 90.1% were classified as species-specific singlets across all modalities. On average, we detected a median of 56 and 57 sgRNA UMIs per cell in HEK293T and NIH/3T3, respectively (**Fig. 1I**). These results indicate that *PerturbFate* achieves a comparable performance in sgRNA capture to previous single-cell CRISPR screening methods(*8*, *16*).

To further evaluate the knockdown effects of the dual-sgRNA system implicated in the *PerturbFate*, we profiled dCas9-KRAB-MeCP2–expressing HEK293 cells transduced with either non-targeting control (NTC) sgRNAs or highly-efficient sgRNAs targeting *DROSHA*, a key regulator of microRNA biogenesis(*8*). We successfully demultiplexed perturbed cells from NTC, indicated by the near-complete loss of chromatin accessibility at the *DROSHA* promoter region (**Fig. 1J**), along with strong suppression of both nascent and pre-existing RNA expression levels of *DROSHA* (88.1% and 86.8% decrease, respectively) in perturbed cells (**Fig. 1K**).

Together, these data establish *PerturbFate* as a novel single-cell platform capable of simultaneously capturing comprehensive molecular modalities with CRISPR perturbations, with key advantages of low cost (less than 0.01$ per cell; **Supplementary Text 1**) and scalability (100,000s to millions of cells per experiment(*3*)). The detailed protocols for *PerturbFate* are included as supplementary files to facilitate individual laboratories to perform high-content CRISPR screenings cost-efficiently.

## Single-cell multi-omics profiling of BRAF^V600E^ melanoma cells via *PerturbFate*

As a demonstration of our method, we applied *PerturbFate* to investigate the core molecular programs mediating Vemurafenib resistance of BRAF^V600E^ melanoma(*22*). Vemurafenib inhibits the hyperactivity of BRAF harboring a *BRAF^V600E^*activating mutation, a key driver in approximately half of all melanoma cases(*22*, *23*). Although Vemurafenib reached a more than 50% response rate, acquired drug resistance and relapse happen in almost all patients(*24*). Although CRISPR screens have identified various potential genetic regulators of Vemurafenib resistance(*25–28*), these regulators function in distinct biological processes (**Fig. S4A**), and the underlying mechanisms linking their primary functions to drug adaptation remain largely unknown. This gap limits our understanding of the molecular basis of Vemurafenib resistance and hinders the development of effective combination therapies for melanoma patients.

A375, a melanoma cell line with a *BRAF^V600E^* mutation that has been used in genome-scale CRISPR-Cas9 screens for identifying genes implicated in Vemurafenib resistance(*27*), was used in the present study. Based on several prior bulk CRISPR screens(*25–28*) and the gene expression profile of A375 cells, we selected 143 candidate genes for targeting (**Fig. 2A, Fig. S4A-B**). A375 cells exhibited sensitivity to Vemurafenib treatment, with 1 μM Vemurafenib showing ∼90% inhibitory effect on cell growth (**Fig. S4C**). Following the construction of an sgRNA plasmid library targeting these drug-resistant candidate genes and subsequent lentiviral packaging, the lentivirus library was transduced into A375 cells with constitutive *dCas9-BFP-KRAB*(*29*) expression at a low MOI (MOI = 0.05) (**Fig. 2A**), and positive cells were sorted after two days of transduction (**Fig. S4D**). After another two days expansion (day 0), cells were then screened in media supplemented with either DMSO or Vemurafenib. Cells for *PerturbFate* profiling were harvested on day 6 to ensure an even sgRNA representation (**Fig. S4E**); cells after 14 days of culture were used for parallel bulk CRISPR screens to reveal sgRNA abundance changes (**Fig. 2A, Fig. S4E**). Our screen showed high consistency across replicates (**Fig. S4F**), and it recapitulated drug-resistant candidate genes that have been identified in previous studies (e.g. *NF1*, *NF2*, *MED12*)(*30–32*) (**Fig. S4G-H**).

**Fig. 2.**
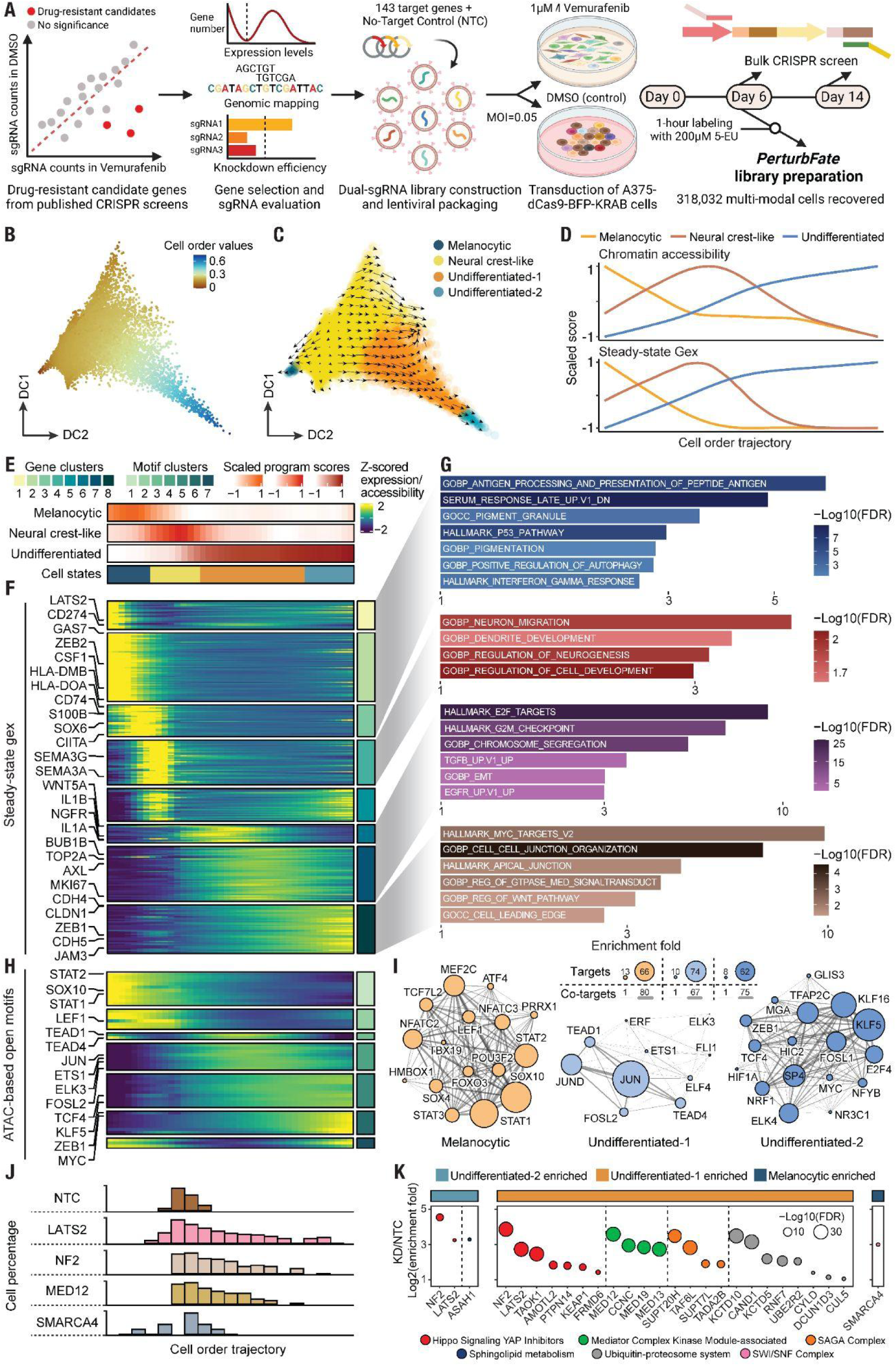
Perturbations of drug-resistant candidate genes significantly drive melanoma phenotype switching toward an undifferentiated state. **(A)** Schematic of the *PerturbFate* experimental design. **(B-C)** Diffusion maps showing the heterogeneous cellular states of genetically perturbed A375 cells under DMSO treatment, colored by their position along the trajectory (**B**) or by annotated cell state (**C**). Arrows in (**C**) indicate the direction of transcriptomic state transitions derived from RNA velocity analysis using nascent and whole-transcriptome data. **(D)** Line plots showing the dynamics of multi-modal melanoma phenotypic gene program scores along the one-dimensional trajectory. The upper panel shows chromatin accessibility scores, computed by averaging normalized counts of ATAC peaks associated with the nascent expression of program member genes. The lower panel shows the steady-state gene expression scores, calculated by averaging the scaled expression values of program member genes in each bin. **(E-H)** Heatmaps with annotation bars showing phenotypic gene program scores and inferred cell states along the trajectory (**E**), scaled steady-state gene expression dynamics (**F**) with representative functional enrichments of state-associated gene programs (**G**), and TF motif accessibility (**H**) along the trajectory. **i,** Network diagrams of state-specific GRNs. Nodes represent TF regulators, with node size corresponding to the number of target genes. Edges indicate shared target genes co-regulated by both TFs, and edge thickness represents the number of co-target genes. **(J)** Histograms showing cell number distributions of representative perturbations identified to induce relative state transitions compared to NTC. *LATS2, NF2, and MED12* perturbations exhibited significant dedifferentiation trends, while *SMARCA4* perturbation induced the differentiation of melanoma cells. **(K)** Dot plots showing the relative enrichment of top perturbations within each cell state compared to NTC. These perturbations also exhibited significant changes in relative cell order distribution compared to NTC. Statistical significance was calculated using proportion tests, and two-sided p-values with FDR correction were visualized by the size of the dots.

Using *PerturbFate*, we obtained multi-omics data from a total of 318,032 cells, including 218,678 cells with linked nascent/pre-existing transcriptomes and chromatin accessibility and 99,354 cells with linked steady-state transcriptomes and chromatin accessibility (**Fig. 2A, Fig. S5A**). On average, 72.4% of cells were successfully matched to sgRNAs, and 71.4% of them were identified as sgRNA singlets (**Fig. S5A-B**). Most perturbations exhibited robust knockdown effects (**Fig. S5C**). As expected, we observed a global improvement in the knockdown efficiency of our dual-sgRNA system compared to our previous single-sgRNA perturbation approach(*8*) (**Fig. S5D**). After filtering out off-target sgRNAs with low efficiencies (**Methods**), we retained 119 genetic perturbations alongside NTC cells, with a median of 479 cells per sgRNA pair in the DMSO condition and 393 cells per sgRNA pair in the Vemurafenib treatment condition. The stable performance of *PerturbFate* was further confirmed across chromatin accessibility, nascent and pre-existing transcriptomes, and whole transcriptomes (**Fig. S5E-G**).

Phenotypic plasticity has been recognized as a key characteristic of melanoma that determines its malignancy(*33*). Using our comprehensive phenotypic profiling, we investigated potential cell state transitions induced by these genetic perturbations. We applied Diffusion Map(*34*) and representative melanoma phenotypic gene signatures(*35*) to project and integrate single-cell transcriptomes onto the diffusion space (**Fig. S6A**), and nascent transcriptome-based RNA velocity was computed to indicate the directionality of cell state transitions. Perturbed A375 cells were ordered into a trajectory encompassing major melanoma phenotypes, including a Melanocytic state, a Neural crest-like state, and an Undifferentiated state, which was further split into two sub-states (**Methods**). Nascent transcriptomes indicate that the primary transition occurs from the Neural crest-like state to the Undifferentiated state, along with another transition from the Neural crest-like to Melanocytic state (**Fig. 2B-C**). Our strategy accurately captures the root of cell state transitions, as original A375 cells have been reported to harbor a Neural crest-like phenotype(*36*). These findings also align with a general association between drug resistance, cancer dedifferentiation, and increased malignancy(*37*). Additionally, by performing ATAC peak-nascent gene expression association analysis(*38*) (**Methods**), we identified cis-regulatory elements (CREs) associated with phenotypic gene signatures and confirmed consistent epigenetic dynamics along the trajectory (**Fig. 2D**).

To dissect functional gene programs linked to these cell states, we examined gene expression patterns along the trajectory (**Methods**). Distinct gene activities underlying different phenotypic cell states were observed: known dedifferentiation markers(*35*) (*e.g., NGFR, WNT5A, AXL*) showed strong activations in the Undifferentiated state, while multiple HLA genes (*e.g., HLA-DMB, HLA-DOA*) and immunogenic markers (*e.g., CD74, CIITA*) exhibited higher expression levels in the Melanocytic state. Also, we observed a strong shift from a *ZEB2*^Hi^ to a *ZEB1*^Hi^ state, a key event in melanocyte dedifferentiation and melanomagenesis(*39*) (**Fig. 2E-F**). Further functional enrichment analysis revealed distinct features of gene clusters associated with each sub-state: cluster 1-3 genes are associated with melanocytic pigmentation and immunogenicity, representing a relatively benign state(*40*); the enrichment of multiple neuronal development terms in cluster 4 genes reflects the Neural crest lineage of steady-state A375 cells. Undifferentiated-1 and Undifferentiated-2 states can be distinguished by cell cycle activity and adhesion regulation, consistent with the slow-cycling property of cancer cells with an undifferentiated mesenchymal phenotype(*33*) (**Fig. 2E-G**). Notably, despite higher noise levels, nascent transcriptomic patterns closely mirrored those from steady-state data (**Fig. S6B**), suggesting that transcriptional regulation, rather than post-transcriptional mechanisms, primarily drives melanoma cell state transitions.

We further examined melanoma cell state-specific epigenetic dynamics along the trajectory by identifying TF motifs with accessibility patterns similar to their gene expression profiles (**Methods**, **Fig. 2H, Fig. S6C**). SOX10, a known master TF driving melanocyte differentiation from the neural crest during the development(*41*, *42*), exhibited the highest motif accessibility in the Melanocytic state (**Fig. 2E-H**). Moreover, the opening of STAT motifs (*e.g.,* STAT1, STAT2) in the Melanocytic state corroborates the activation of the Interferon (IFN) response and antigen presentation observed at the gene expression level (**Fig. 2E-H**). In contrast, the transition from the Neural Crest-like state to the Undifferentiated state was accompanied by increased motif accessibility of TEADs (*e.g.,* TEAD1, TEAD4) and AP-1 family motifs (*e.g.,* JUN, FOSL2) (**Fig. 2E-H**), consistent with prior findings that stepwise SOX10 loss and AP-1 and TEAD activation epigenetically reprograms melanoma cells and promotes Vemurafenib resistance(*2*).

To obtain an integrative view of multi-modal temporal dynamics, and to further elucidate the gene regulatory principles driving the melanoma state transition, we used an analytical strategy that leverages our single-cell multimodal data to map TF-gene regulatory linkages (**Fig. S6D**): First, we identified candidate TF-target gene pairs based on the association between nascent gene expression and chromatin accessibility of nearby ATAC peaks, and the embedding of TF motifs within gene associated peaks. Following the basic principle of transcriptional activation where TF binding opens chromatin regions of CREs, leading to RNA synthesis activation and ultimately steady-state gene upregulation, we then refined these linkages using temporal constraints. Only TF-target gene pairs in which TF motifs, nascent gene expression, and steady-state gene expression peaked following this order along the state transition were retained (**Methods**). We reconstructed three major GRNs driving transitions toward the melanocytic, undifferentiated-1, and undifferentiated-2 states, respectively (**Fig. 2I**). For example, despite its increased motif accessibility, SOX10 regulon (*i.e.,* a group of genes co-regulated by SOX10) emerged as the dominant regulon in the melanocytic GRN(*41*). MEF2C, another TF, was reported to collaborate with SOX10 to direct neural crest toward a melanocytic fate(*43*). Their cooperation was also revealed by a substantial overlap between their target genes: 53 of 72 MEF2C targets (73.6%) and 104 SOX10 targets (51.0%) were shared (**Fig. S6E-G**).

By identifying TF regulons that contribute to transitions toward undifferentiated cell states, we observed that AP-1 family factors (*e.g.,* JUN, JUND, FOSL2) downstream of MAPK signaling and TEAD TFs (*e.g.,* TEAD1, TEAD4) downstream of Hippo/YAP signaling are major regulons (**Fig. 2I**), corroborating previous studies highlighting both AP-1 and TEADs as critical mediators of melanoma dedifferentiation(*44*, *45*). Additionally, substantial target gene overlap between AP-1 and TEADs regulons in the Undifferentiated-1 GRN (**Fig. S6F-G**) further supports their established synergistic roles in driving tumor invasion and metastasis(*46*, *47*). The Undifferentiated-2 state GRN is characterized by regulons of multiple TFs known to participate in tumor progression and metastasis, including KLF5, KLF16, ZEB1, MYC, and TCF4(*48–51*), with FOSL1 as the sole AP-1 factor (**Fig. 2I**). This is consistent with a previous study identifying a more concentrated activation of FOSL1 than JUN and FOSL2 in the undifferentiated melanoma state(*52*), (*^53^*).

To quantify the effects of individual genetic perturbations on cell state transition, we compared the cell order distribution between perturbations and NTC and conducted a cell state-specific enrichment analysis of perturbations relative to NTC (**Methods**, **Fig. S6H**). As expected, 52 perturbations significantly altered the cell state distribution within the trajectory. The majority (46 of 52) were relatively more undifferentiated, and 33 of these 46 perturbations also showed significant enrichment in the Undifferentiated states (**Figure 2J-K**, **Fig. S6I**). Only the *SMARCA4* perturbation induced both a significant shift toward the Melanocytic state and significant enrichment in the Melanocytic state (**Fig. 2J-K**). We noticed that target genes of perturbations inducing melanoma dedifferentiation converged to similar biological functions. For example, perturbations targeting genes encoding Hippo signaling YAP inhibitors (*e.g., NF2*, *LATS2*, *PTPN14*, *TAOK1*) exhibited the strongest shift toward an undifferentiated state (**Fig. 2K**), closely matching the activation of TEADs regulons revealed by our GRN analysis. Moreover, perturbations targeting genes encoding components of the Mediator complex (e.g., *MED12*, *MED13*, *CCNC*, *MED19*), the SAGA complex (*e.g., SUPT20H*, *TAF6L*, *TADA2B*), and the ubiquitin-proteasome system (*e.g., KCTD10*, *CAND1*, *RNF7*, *KEAP1*) also induced robust melanoma dedifferentiation (**Fig. 2K**). Multi-modal regulon scoring by integrating gene expression, motif accessibility, and nascent expression of target genes (**Methods**) confirmed a general activation of AP-1 and TEADs regulons across perturbed cell populations that received sgRNAs targeting genes spanning diverse functional categories **Fig. S6J**), suggesting the transition of melanoma to an undifferentiated state is largely mediated through convergent molecular programs.

## Identification of convergent gene regulatory programs driving Vemurafenib resistance

To investigate how Vemurafenib treatment alters the cell state dynamics of melanoma cells, we projected both Vemurafenib-and DMSO-treated cells onto the same trajectory (**Fig. 3A-B**), following a similar approach as that used for DMSO-treated cells (**Fig. 2B**). Consistent with our findings linking AP-1 activation to melanoma dedifferentiation (**Fig. 2I**), Vemurafenib, a BRAF-MAPK signaling inhibitor, induced a widespread shift toward a melanocytic state in melanoma cells, as confirmed by nascent transcriptome-based RNA velocity analysis and phenotypic signature scoring (**Fig. 3B-D**).

**Fig 3.**
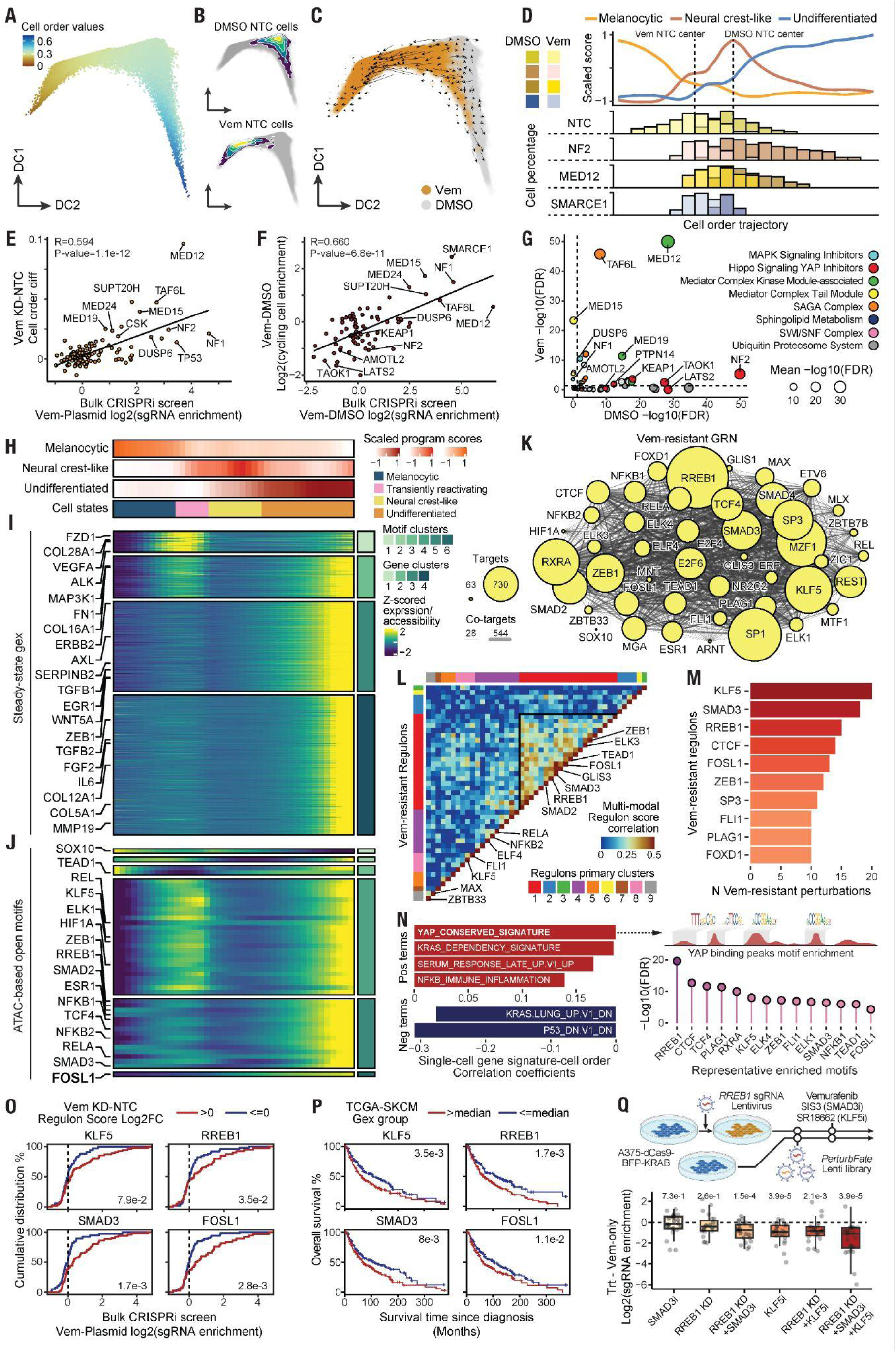
Identification of the molecular programs driving drug resistance in melanoma cells. (A-C) Diffusion maps showing the heterogeneous cell states of genetically perturbed A375 cells under DMSO and Vem treatment. In **(A)**, cells are colored by their “order” along the trajectory; in **(B)**, densities of NTC cells under DMSO and Vemurafenib treatment conditions are highlighted; in **(C)**, they are colored by treatment condition, with arrows indicating transition directions derived from nascent transcriptome-based RNA velocity. **(D)** Line plot showing steady-state transcriptome-based melanoma phenotypic gene program scores along the integrated trajectory that includes both DMSO- and Vem-treated cells (upper panel). The lower panel shows histograms displaying the cell-state distribution of representative perturbations along the trajectory, grouped by treatment condition. **(E)** Scatter plot comparing relative cell state shift of genetic perturbations to NTC with growth advantage of perturbations over NTC revealed by bulk CRISPR screen. **(F)** Scatter plot comparing the relative cell cycle activity change of perturbations upon Vemurafenib treatment and growth phenotype change of perturbations upon Vemurafenib treatment. Cell cycle activity was represented by the proportion of cells computationally assigned to be at S/G2M phases under each perturbation. **(G)** Scatter plot comparing the significance of the relative dedifferentiation shift of each perturbation to NTC under DMSO and Vemurafenib treatment conditions. **(H)** Heatmaps showing phenotypic gene program scores and inferred cell states along the trajectory containing only Vemurafenib-treated cells. **(I-J)** Heatmaps illustrating the scaled gene expression dynamics (**i**) and TF motif accessibility (**j**) along the trajectory. **(K)** Network diagram of Vemurafenib-resistant GRN. Each TF node’s size indicates the number of target genes; edges connect TFs that share co-targets, with line thickness corresponding to the number of shared targets. **(L)** Heatmap showing correlations of multi-modal regulon scores among Vem-resistant regulons under Vemurafenib treatment; hierarchical clustering identified groups of co-varying regulons, with cluster1 exhibiting the highest degree of covariation pattern. **(M)** Bar chart ranking the top 10 Vem-resistant regulons by the number of perturbations that significantly elevate their scores compared to NTC cells under Vemurafenib treatment. **(N)** Bar chart showing correlations between single-cell Oncogenic Signatures and cell order values. Top terms exhibiting significant correlations with cell orders (by a permutation test, FDR-corrected) are shown (left panel). The right panel shows representative TF motifs that exhibit strong local enrichment in YAP ChIP-seq peaks and their enrichment FDR-corrected two-sided p-values. **(O)** Line plots showing the cumulative distribution of relative growth advantages of perturbations, which are stratified by their relative multi-modal regulon score changes compared to NTC. K-S tests were conducted to examine the statistical significance of the distribution difference. **(P)** Kaplan-Meier survival curve plots comparing overall survival of patients in the TCGA SKCM cohort. Patients were stratified by the expression levels of key TFs. Statistical significance was examined by log-rank tests. **(Q)** Validation of anti-Vemurafenib resistance effects of combined inhibition of candidate TFs. The upper panel illustrates the experimental design, in which additional genetic and/or chemical perturbations were conducted in parallel with the *PerturbFate* bulk CRISPR screen. The lower panel shows the box plot demonstrating the gradually improved efficacy on inhibiting growth advantage of drug-resistant perturbations (n=24) by combining key TFs identified to drive the transition to a Vemurafenib-resistant Undifferentiated state. FDR-corrected two-sided p-values represent the significance of the global deviation from 0.

We evaluated how various genetic perturbations influence the cellular response to Vemurafenib treatment (**Fig. S7A**). Compared to the NTC, perturbations exhibited resistance to Vemurafenib-induced melanoma differentiation on day 6 to varying extents, which significantly correlated with their growth advantage over the NTC on day 14 estimated from the bulk CRISPR screen (pearson correlation coefficient r = 0.594, p-value = 1.1e-12; **Fig. 3E**, **Fig. S7A-B**). Furthermore, cell cycle activity inferred from *PerturbFate* single-cell gene expression also aligned with their growth phenotypes (pearson correlation coefficient r = 0.66, p-value = 6.8e-11; **Fig. 3F**), reinforcing the strong association between melanoma cell state plasticity and cell growth of drug-resistant perturbations.

We identified 24 perturbations, such as *NF2*, *MED12*, and *SMARCE1*, that exhibited resistance to both growth arrest, as measured by the bulk CRISPR screen (**Fig. 4H**), and to cellular differentiation, as measured by *PerturbFate*, relative to NTC cells under Vemurafenib treatment (**Fig. 3D-F**). Most of these Vemurafenib-resistant perturbations (20 out of 24) also induced significant melanoma dedifferentiation in the DMSO condition, although the extent of dedifferentiation varied between the two conditions (**Fig. 3G**). For example, perturbations targeting inhibitors of the Hippo signaling (e.g., *NF2*, *TAOK1*, *KEAP1*, *PTPN14*) strongly induced an undifferentiated state in melanoma cells under DMSO but had dramatically reduced effects under Vemurafenib treatment (**Fig. 3G**). This aligns with previous studies identifying the MAPK pathway as a key promoter of the downstream effects of Hippo signaling in the context of cell proliferation, drug treatment, and cancer resistance(*58–60*). In contrast, perturbations targeting MAPK signaling inhibitors (e.g., *NF1*, *DUSP6*) had little effect in DMSO yet promoted cell proliferation and dedifferentiation under Vemurafenib treatment (**Fig. 3G**). Meanwhile, some perturbations, such as those targeting genes encoding components of the Mediator Complex (e.g., *MED12*, *MED19*), consistently promoted cellular dedifferentiation under both conditions (**Fig. 3G**).

**Fig. 4.**
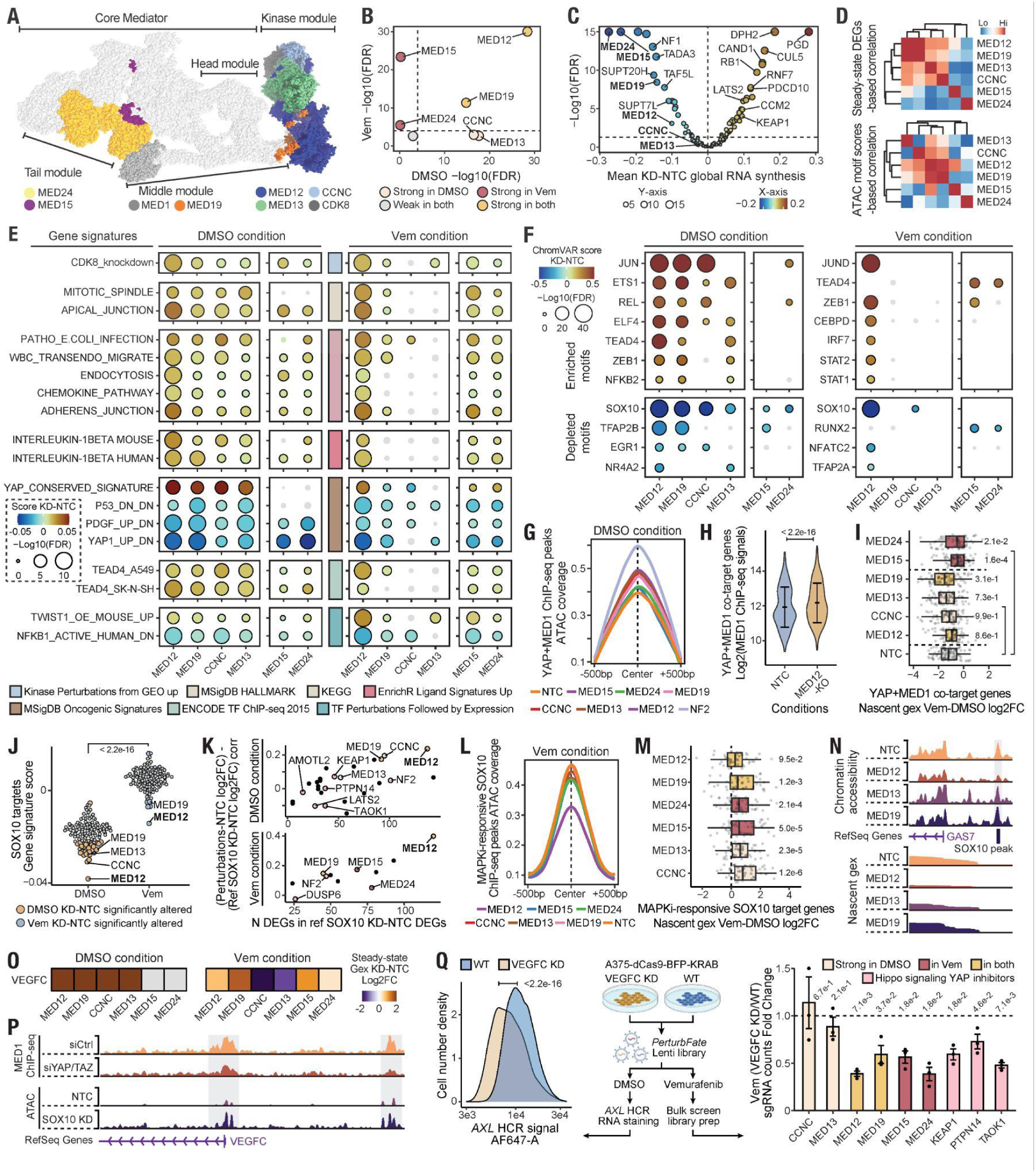
Perturbations of distinct Mediator Complex components drive drug resistance through divergent mechanisms. **(A)** Structural diagram of the human Mediator Complex, highlighting the components analyzed and discussed in this study. The structure is based on published cryo-electron microscopy data(*80*). **(B)** Scatter plot showing the statistical significance of relative dedifferentiation shifts of cells with Mediator Complex components perturbed compared to NTC cells under DMSO (x-axis) and Vemurafenib (y-axis) treatment conditions. **(C)** Scatter plot illustrating the effects of genetic perturbations on global RNA transcription changes compared to NTC cells. Global RNA synthesis is represented as log-transformed, aggregated nascent gene expression for the top 1,000 genes in NTC cells. **(D)** Heatmaps with hierarchical clustering depicting transcriptomic (upper panel) and epigenetic motif accessibility (lower panel) similarities across Mediator Complex component perturbations in the DMSO condition. **(E-F)** Dot heatmaps showing the upregulation/downregulation of phenotypic gene signatures (**E**) and the enrichment/depletion of TF motifs (**F**) across Mediator Complex component perturbations, along with statistical significance. Dot color represents relative upregulation/downregulation for gene signatures or enrichment/depletion for TF motifs. Dot size indicates statistical significance, and gray dots denote non-significance. **(G)** Line plot comparing ATAC signal coverage at YAP-Core-Mediator co-regulatory regions upon Mediator Complex component perturbations in the DMSO condition. NTC and *NF2*-perturbed cells are shown as negative and positive controls, respectively. YAP-Core Mediator co-regulatory regions are defined as YAP ChIP-seq peaks overlapping MED1 ChIP-seq peaks, which are absent in the siYAP/TAZ condition (reprocessed from(*74*)) (**Fig. S9G**). **(H)** Violin plot displaying the signal distribution of MED1 ChIP-seq peaks within YAP-Core Mediator co-target genes (n=295) between NTC and *MED12* knockout (KO) conditions. To examine the direct inhibitory effects of the Kinase Module on the transcriptional activity of Core-Mediator, only MED1 ChIP-seq peaks overlapping with MED12 ChIP-seq peaks were considered (reprocessed from(*72*)). A paired Wilcoxon test was conducted, and a two-sided p-value is shown. **(I)** Box plot illustrating global nascent gene expression changes for YAP-Core Mediator co-target genes (n=75) across Mediator Complex component perturbations following Vemurafenib treatment. One-way ANOVA followed by Tukey’s post hoc tests was performed, with FDR-corrected two-sided p-values displayed for comparisons between perturbations and NTC cells. **(J)** Beeswarm plot showing the distribution of averaged perturbation-specific SOX10 target gene signature(*75*) scores in DMSO and Vemurafenib treatment conditions (n=120). A paired Wilcoxon test was performed, and a two-sided p-value is shown. Perturbations with significant distribution changes in single-cell signature scores compared to NTC cells are colored in orange (DMSO condition) and purple (Vemurafenib condition). **(K)** Scatter plots showing the overlap between strong perturbations-induced DEGs and SOX10 KD-induced DEGs (reprocessed from (*75*)) (x-axis), as well as the correlation of expression changes in *SOX10* KD-induced DEGs between strong perturbations and *SOX10* KD. DMSO (upper panel) and Vemurafenib (lower panel) treatment conditions are plotted separately. **(L)** Line plot comparing ATAC signal coverage at MAPKi-responsive SOX10 binding regions (reprocessed from(*76*), **Fig. S10E**) upon Mediator Complex component perturbations in the Vemurafenib treatment condition. NTC cells are shown as controls. **(M)** Box plot displaying global nascent gene expression changes for MAPKi-responsive SOX10 target genes (reprocessed from(*76*), **Fig. S10E**) (n=46) across Mediator Complex component perturbations following Vemurafenib treatment. One-sample Wilcoxon tests comparing the deviation of each sample from 0 were performed, and FDR-corrected two-sided p-values are shown. **(N)** Representative genome tracks illustrating impaired SOX10 proximal peak and promoter chromatin accessibility (upper panel) and nascent transcription (lower panel) of *GAS7*, a melanocytic gene, upon *MED12* perturbation compared to other Kinase Module-associated Mediator Complex perturbations. **(O)** Heatmaps showing expression change patterns across Mediator Complex component perturbations compared to NTC in DMSO and Vemurafenib treatment conditions. Gray tiles represent a maximum CPM < 20 in the comparison, indicating low expression. **(P)** Genome tracks demonstrating that the transcriptional activity of *VEGFC* is dependent on Hippo/YAP (reprocessed from(*74*)) and that its promoter chromatin accessibility is activated upon SOX10 loss-of-function (reprocessed from(*75*)). **(Q)** Validations of *VEGFC* as a key convergent factor inducing Mediator Complex component perturbation-driven melanoma dedifferentiation and drug resistance. VEGFC-KD dCas9-expressing A375 cells were transduced with the *PerturbFate* sgRNA library to induce dedifferentiation and drug resistance. *AXL* HCR staining followed by flow cytometry was performed to compare the global undifferentiated state of cells in the DMSO condition. A K-S test was conducted, and a two-sided p-value is reported. The right panel shows relative growth reductions of key perturbations upon additional *VEGFC* KD under Vemurafenib treatment (n=3). One-sample Wilcoxon tests were performed to examine deviation from 1, and FDR-corrected two-sided p-values are displayed. Boxes in box plots indicate the median and interquartile range (IQR), with whiskers indicating 1.5× IQR.

We then examined the multi-modal molecular dynamics in Vemurafenib-treated cells. Similar to cells in the DMSO condition, we observed continuous alterations in phenotypic gene programs and extensive epigenetic remodeling, enabling us to classify Vemurafenib-treated cells into melanocytic, neural crest-like, and undifferentiated states (**Fig. 3H-J, Fig. S7C-D**). These cellular states were further validated by elevated SOX10 activity in the differentiated state and FOSL1 motif accessibility in the undifferentiated state (**Fig. 3H-J**). Notably, based on coordinated transitions in global chromatin accessibility, open peak features, and nascent transcription, we further identified a transiently reactivating state in Vemurafenib-treated cells during their transition toward the differentiated melanocytic state (**Fig. 3H**, **Fig. 7C-J**). Aligning with a sharp wave of global promoter opening (**Fig. 3H-I**), we observed the partial activation of undifferentiated state-associated TF motif accessibility and gene expression within this state (**Fig. 3H-J**). However, the lack of activities of MAPK-downstream serum response factors (ELK1, ELK3, ELK4), RREB1, and especially FOSL1, may have impeded the further activation of undifferentiated state-associated gene regulation (**Fig. S7K-M**).

By reconstructing GRNs for transitions of Vemurafenib-treated cells to the Undifferentiated state using our established strategy (**Methods, Fig. 6D**), we uncovered a dense network of TF regulators that collaboratively drive the transition toward the Vemurafenib-resistant Undifferentiated state (**Fig. 3K**), and further confirmed its partial activation in the Transiently reactivating state (**Fig. 8A-D**). Notably, this Vemurafenib-resistant undifferentiated state is characterized by the concurrent activation of FOSL1 downstream of MAPK signaling, TEAD1 downstream of Hippo signaling, SMAD2/3/4 downstream of TGF-β signaling, TCF4 downstream of canonical Wnt signaling, the epithelial-mesenchymal transition (EMT)-associated TF ZEB1, and additional TFs less characterized in melanoma dedifferentiation and drug resistance, such as RREB1 and KLF5 (**Fig. 3K**).

To further dissect the potential roles of these TF in drug resistance, we quantified their multi-modal regulon activities within each perturbation (**Methods**). By analyzing pairwise correlations between TF regulons across perturbations (**Methods**), we identified a cluster of regulons that exhibited substantial co-variation across perturbations (**Fig. 3L**), potentially representing a synergistic regulatory hub underlying drug resistance. In addition to FOSL1 and TEAD1, ELK3 and ZEB1, which are known to co-promote epithelial-to-mesenchymal transition (EMT)(*61*, *62*), were clustered closely. Moreover, we observed strong co-variations between SMADs and RREB1, which has been recently reported to co-activate EMT in both pancreatic cancer(*63*) and KRAS-activated lung cancer(*64*).

Furthermore, by examining the significance of the activation of these TF regulons across 24 drug-resistant perturbations (**Methods**), we found that KLF5, SMAD3, RREB1, and FOSL1 were the most broadly activated regulons, which might represent key convergent gene programs (**Fig. 3M**). Although the KLF5 regulon grouped with FLI1 (**Fig. 3L**), TEAD1 and FOSL1 emerged as another two of its most co-varying partners. The existence of KLF5, SMAD3, and FOSL1 motifs within accessible peaks associated with RREB1 implies a potential direct regulatory relationship among these TFs (**Fig. S8E**).

By correlating single-cell oncogenic signatures and their orders in the trajectory (**Methods**), we found that the YAP signature, along with the KRAS dependency signature, was most positively associated with Vemurafenib-resistant dedifferentiation (**Fig. 3N**). This finding is in line with both of our perturbation-level (**Fig. 3G**) and molecular-level examinations (**Fig. 3I-J**), underscoring the essential role of both the Hippo/YAP and MAPK pathways during melanoma drug adaptation. Again, several key transcription factors, including RREB1, KLF5, SMAD3, and FOSL1, were found to be strongly enriched in YAP genomic binding peaks, as revealed by external YAP ChIP-seq data(*65*). Broad co-dependencies among these genes, as captured by DepMap(*66*), further reinforce their functional convergence (**Fig. S8G**). Moreover, when we stratified perturbations based on the relative regulon score changes of RREB1, KLF5, SMAD3, and FOSL1 compared to NTC, we observed that activation of these regulons predisposed cells to significantly enhanced Vemurafenib resistance in our bulk screen (**Fig. 3O**). The strong negative association between the expression of these TFs and worse patient prognosis in the TCGA Skin Cutaneous Melanoma (SKCM) cohort(*67*) (**Fig. 3P**) further underscores the essential role of these TFs and their associated gene programs in mediating melanoma malignant transformation.

Finally, we sought to validate the collaborative roles of these less-characterized TFs in mediating Vemurafenib resistance. Specifically, we performed secondary CRISPR screens under Vemurafenib treatment, incorporating additional genetic or chemical perturbations targeting KLF5, RREB1, and SMAD3 (**Methods**). Across 24 drug-resistant perturbations, while inhibition of SMAD3 or RREB1 alone did not significantly decrease overall drug resistance, their combined inhibition resulted in a substantial reduction in global growth advantage (**Fig. 3Q**). In line with our earlier identification of KLF5 as a critical regulator of melanoma dedifferentiation under both DMSO and Vemurafenib treatment conditions (**Fig. 2I**, **3K**), inhibition of KLF5 alone was sufficient to significantly reverse Vemurafenib resistance (**Fig. 3Q**). Moreover, the combined inhibition of RREB1, SMAD3, and KLF5 produced the most pronounced broad-spectrum reduction in drug resistance across heterogeneous perturbations, with a mean 3.1-fold reduction in relative growth advantage compared to the Vemurafenib-only condition (**Fig. 3Q**), confirming their synergies in driving Vemurafenib resistance in melanoma cells.

## Perturbation of Mediator Complex genes results in structure-concordant, heterogeneous melanoma cell state transitions

We next investigated the molecular mechanisms by which different genetic perturbations influence cell state dynamics under both DMSO and Vemurafenib treatments, with a particular focus on the Mediator Complex, a crucial regulator of transcription initiation, pausing, and elongation(*70*). Several components of this complex are implicated in promoting melanoma dedifferentiation and resistance to Vemurafenib treatment (**Fig. 2K, 3G**), although molecular events underlying these functional associations remain elusive.

The Mediator Complex is organized into multiple modules: the Head, Middle, Tail, and Kinase Modules(*71*) (**Fig. 4A**). The Kinase Module, comprising MED12, MED13, CCNC, and CDK8, functions by sterically hindering RNA Polymerase II from binding to the Core Mediator, a key component of the transcriptional Pre-Initiation Complex (PIC) required for transcriptional activation(*70*). In contrast, the Core Mediator, formed by the Head, Middle, and Tail modules, facilitates transcription by bridging transcription factors bound to the Tail module with the PIC. Interestingly, perturbations of different Mediator Complex components resulted in heterogeneous responses corresponding to their structural compositions (**Fig. 4A-B, Fig. S9A**): under the DMSO condition, knockdown of Kinase Module genes (*i.e., MED12, MED13, and CCNC*), and *MED19*, which encodes a Middle Module component extensively interacting with MED12, induced strong melanoma dedifferentiation (**Fig. 4B**). In contrast, knockdown of Tail Module genes (*i.e., MED15, MED24*) led to dedifferentiation and drug resistance under Vemurafenib treatment but failed to induce state transitions under the DMSO condition (**Fig. 4B**). Furthermore, nascent transcriptome-based global RNA synthesis analysis revealed that *MED15* and *MED24* perturbations caused the most significant transcriptional impairment (**Fig. 4C**), whereas Kinase Module-associated perturbations (*i.e., MED12, MED19, MED13, CCNC*) exhibited highly similar transcriptomic and epigenetic landscapes (**Fig. 4D**). UMAP embeddings of perturbed transcriptomes and chromatin landscapes further confirmed their functional discrepancies (**Fig. S9B-C**). These observations suggest that module-specific biochemical functionalities contribute to distinct perturbed cellular phenotypes.

To identify convergent molecular programs contributing to intra-module similarity and inter-module diversity, we performed perturbation-specific transcriptomic phenotyping and motif activity analysis (**Methods**), revealing various gene signatures and TF motifs that showed consistent changes across Kinase Module-associated perturbations under the DMSO condition (**Fig. 4E-F, Fig. S9E-F**). For instance, the activated apical junction signature aligns with the undifferentiated states of these perturbed cells (**Fig. 4E**). Furthermore, the activation of inflammatory gene programs (*e.g.,* IL1β, chemokine signaling, and NFKB1 activation), coupled with increased AP-1 and NFKB motif activities (**Fig. 4E-F, Fig. S9E-F**), supports the role of the Kinase Module in suppressing inflammation via direct suppression of the Core-Mediator activity(*72*) or the recruitment of chromatin repressors(*73*). Among these gene signatures, Hippo/YAP signaling-related terms and TF motifs (*e.g.,* YAP, TEAD4) exhibited the strongest activation following the perturbation of the Kinase Module (**Fig. 4E-F**). This functional association is supported by UMAP embeddings, where Kinase Module perturbations cluster near Hippo signaling YAP inhibitor perturbations (**Fig. S9B-C**), and by the co-dependencies between Kinase Module genes and Hippo signaling YAP inhibitor genes captured by DepMap(*66*) (**Fig. S9D**). To validate the potential role for the Kinase Module in regulating the Hippo/YAP-mediated gene program, we analyzed external ChIP-seq data(*74*), and revealed a strong overlap between the genomic binding regions of the Core Mediator and YAP (**Fig. S9G**). We further identified Core-Mediator binding regions with functional and physical dependence on YAP and their associated target genes (**Fig. S9G**), and observed increased chromatin accessibility at these sites upon the Kinase Module perturbation (**Fig. 4G**). Additionally, within these target genes, such as *VEGFC*, *TGFBR2*, and *GLIS3* (**Fig. S9H**), we used another set of external ChIP-seq data(*72*) to confirm that *MED12* knockout led to a significantly increased Core-Mediator binding signal at MED1 peaks that overlap with MED12 peaks (**Fig. 4H**). Given the critical role of Hippo/YAP signaling in driving melanoma dedifferentiation, these findings suggest that the Kinase Module suppresses melanoma dedifferentiation by directly inhibiting the transcriptional activity of the Core Mediator at YAP regulatory elements.

In contrast, perturbations in the Tail Module (*MED15*, *MED24*) had no detectable impact on melanoma dedifferentiation or Hippo/YAP signaling activation in the DMSO condition (**Fig. 4A-B**). However, following Vemurafenib treatment, these gene signatures remained significantly elevated in *MED15*- and *MED24*-perturbed cells compared to NTC controls, unlike some Kinase Module perturbations (*CCNC* and *MED13*) (**Fig. 4E**). Consistent with the severely impaired transcriptional control upon Tail Module knockdown (**Fig. 4C**), we observed pronounced resistance of *MED15*- and *MED24*-perturbed cells to Vemurafenib-induced differentiation, along with a less altered global chromatin landscape, implying their lack of response to Vemurafenib treatment (**Fig. S9I-J**). Moreover, nascent gene expression analysis confirmed the less inhibited transcription of YAP-Core-Mediator co-target genes in *MED15* and *MED24* perturbations upon Vemurafenib treatment (**Fig. 4I**). The elevated gene signatures and increased motif accessibility of TEAD4 in *MED15*- and *MED24*-perturbed cells further supported our results, highlighting the critical role of Hippo/YAP signaling in driving the dedifferentiation of melanoma cells upon Vemurafenib treatment.

Unlike perturbations of genes encoding other components of the Mediator Complex, perturbing *MED12* profoundly promoted the dedifferentiation of melanoma cells under both DMSO and Vemurafenib treatments and exhibited the highest resistance to Vemurafenib-induced differentiation (**Fig. 4B, Fig. S9I**). This suggests that *MED12* knockdown may drive melanoma dedifferentiation and drug resistance through mechanisms independent of the Kinase Module. Specifically, *MED12* knockdown led to the most dramatic depletion of SOX10 activity at both the epigenetic and transcriptomic levels. This depletion persisted even after Vemurafenib treatment, which normally increases SOX10 activity and directs the differentiation of melanoma cells (**Fig. 4F, 4J, Fig. S10A**). Given SOX10’s essential role in directing melanocytic cell fate (**Figure 2i**) and the fact that its knockdown is sufficient to induce an undifferentiated melanoma state with restored AP-1 activity(*75*), we hypothesized that SOX10 loss-of-function contributes to the effects of *MED12* knockdown. The unique similarity of transcriptomic responses between *MED12*-perturbed cells and reference *SOX10*-perturbed melanoma cells(*75*) further supports this hypothesis (**Fig. 4K, Fig. S10B-D**). Using paired external ChIP-seq and RNA-seq data(*76*) (**Fig. S10E**), we further compared the strength of SOX10-mediated effects upon Vemurafenib treatment across different Mediator Complex component perturbations. *MED12* knockdown impaired the chromatin accessibility of MAPK inhibitor (MAPKi)-induced SOX10 binding regions (**Fig. 4L**) and reduced the activation of MAPKi-induced SOX10 target genes (**Fig. 4M**). For example, the chromatin accessibility of a validated proximal SOX10 binding site regulating *GAS7*, a melanocytic gene(*77*) (**Fig. 2F**), and its nascent transcription were both strongly affected only in *MED12* knockdown cells (**Fig. 4N**). These findings suggest a unique loss of SOX10 function in mediating melanoma dedifferentiation upon *MED12* perturbation, aligning with previous reports on the role of MED12-SOX10 interactions in driving the differentiation of neural crest-derived myelinating glial cells(*78*).

In addition to identifying convergent gene programs, we investigated key convergent genes that promote dedifferentiation and Vemurafenib resistance across Mediator Complex component perturbations. Given the established role of the Kinase Module in suppressing cytokine expression, we analyzed cytokine expression patterns across Mediator Complex component perturbations. We observed that only the activation of *VEGFC*, which has been identified as a Vemurafenib resistance marker in melanoma(*2*) and whose autocrine signaling has been implicated in cancer development(*79*), corroborated dedifferentiation patterns under both treatment conditions (**Fig. 4O**, **Fig. S9K**). Interestingly, we identified *VEGFC* as a YAP-Core-Mediator co-regulatory target gene (**Fig. 4P**, **Fig. S9H**), and its expression showed a strong positive correlation with other Hippo/YAP signaling target genes in the TCGA SKCM cohort (**Fig. S9L**). Its drastic activation following direct *SOX10* knockdown further suggests that it may serve as a core downstream mediator of melanoma dedifferentiation (**Fig. 4P**, **Fig. S10C**).

We generated A375 cells with *VEGFC* knockdown and conducted secondary CRISPRi bulk screens (**Fig. 4Q**). *VEGFC* knockdown led to a global downregulation of *AXL*, a marker of the undifferentiated melanoma state (**Fig. 4Q**), and suppressed the overall growth of perturbed melanoma cells under Vemurafenib treatment (**Fig. S10F**). Consistent with our analyses, *VEGFC* knockdown significantly reduced the growth advantage of *MED12*, *MED19*, *MED15*, and *MED24* perturbations, whereas *CCNC-* and *MED13*-perturbed cells remained unaffected (**Fig. 4Q**, **Fig. S10G**). Additionally, *VEGFC* knockdown also suppressed the growth of Hippo signaling YAP inhibitors perturbations (**Fig. 4Q**), further supporting its role in Hippo/YAP-mediated melanoma dedifferentiation.

In summary, our results demonstrate that *PerturbFate* enables the identification of shared cellular states and molecular features driving cancer drug resistance across various perturbations through massively parallel multi-modal profiling. Additionally, it facilitates the in-depth dissection of perturbation-specific effects, as exemplified by Mediator Complex component perturbations. These unique features highlight *PerturbFate* as a powerful tool for uncovering both common and distinct regulatory mechanisms contributing to therapeutic resistance in cancer cells.

## Discussion

Here we introduce *PerturbFate*, a highly scalable single-cell multi-omics screening platform. *PerturbFate* represents the first technique that enables the investigation of the effects of large-scale genetic perturbations on the continuum of gene regulation, from chromatin remodeling to nascent transcription and ultimately to steady-state transcriptomic phenotypes, at single-cell resolution. By integrating the temporal dynamics of all three molecular layers, *PerturbFate* facilitates the reconstruction of intricate gene regulatory networks by mapping transcription factor modules based on their expression levels, motif accessibility, and target gene transcription, thereby providing deeper insights into the critical drivers of cell state transitions. Importantly, *PerturbFate* is built on an optimized single-cell combinatorial indexing strategy(*8*), significantly enhancing throughput (handling millions of cells) and reducing library preparation costs to less than $0.01 per cell, orders of magnitude lower than conventional methods. This high-throughput capability provides unique opportunities to uncover both shared and unique gene regulatory networks across a vast array of genetic perturbations.

To demonstrate the unique advantage of *PerturbFate* in dissecting complex cell fate dynamics, we applied it to a BRAF^V600E^ melanoma model, focusing on genetic regulators implicated in Vemurafenib resistance. We captured transitions from a Neural crest–like state toward either a differentiated (Melanocytic) or Undifferentiated melanoma state, which mimics the phenotypic plasticity of melanoma in vivo and in vitro(*33*). Through integrated analyses of chromatin accessibility and both steady-state and nascent transcriptomes, we identified key TF modules governing these transitions. Notably, SOX10 serves as the major regulator of the melanocytic state, whereas AP-1 factors and the TEADs factors emerged as key drivers of dedifferentiation.

Vemurafenib treatment that inhibits the BRAF-MAPK pathway induces strong cellular differentiation toward the Melanocytic state. However, perturbations derepressing MAPK pathway or YAP activity consistently directed cells toward a more invasive dedifferentiated state, underscoring these pathways as critical therapeutic targets. Moreover, drug-resistant perturbations, despite that these target genes encoding proteins with distinct functions, circumvented the drug-induced growth defect by reprogramming cellular chromatin landscape into a shared Vemurafenib-resistant molecular state. In addition to well-characterized AP-1 and TEADs, we further identified TFs such as KLF5, RREB1 and SMAD3, whose activities were strongly linked to this resistant state across diverse perturbations. Notably, targeted perturbations of these shared TF modules neutralized the effects of most drug-resistant perturbations, indicating that these factors act as central nodes within a common pro-resistance network.

Although drug-resistant cells induced by distinct drug-resistant perturbations converge on a similar molecular state, the mechanisms underlying these genetic perturbation-to-state linkages remain largely unknown. One striking example involves perturbations affecting various modules of the Mediator complex. The knockdown of Kinase Module-associated genes (*MED12, MED13, CCNC, MED19*) primarily promotes dedifferentiation under DMSO conditions, in part by unblocking the transcriptional activity of the YAP-mediated gene program and the expression of inflammatory genes. In contrast, perturbations of the Tail Module (*MED15, MED24*) induce dedifferentiation exclusively under Vemurafenib treatment by preserving YAP target gene transcription. Furthermore, *MED12* knockdown specifically promotes a strong dedifferentiation trend in melanoma cells under both DMSO and Vemurafenib conditions, which is associated with the loss of SOX10 activity. Our analyses reveal the functional heterogeneity among Mediator complex components in reprogramming melanoma phenotypic states, achieved through both complex-dependent and independent mechanisms. Meanwhile, the unified activation of the YAP-driven gene program, in which *VEGFC* plays a pivotal role, underscores the essential role of this convergent program in driving a more dedifferentiated cell state.

In conclusion, our study establishes *PerturbFate* as a transformative platform for dissecting multiple layers of gene regulation at single-cell resolution. By integrating large-scale single-cell multi-modal readouts, *PerturbFate* provides insights into key regulators of cell fate decisions. Beyond melanoma and Vemurafenib resistance, *PerturbFate* provides a foundational framework for understanding the development and transition of disease-associated cellular phenotypes.

## Methods

### Cell culture

The A375 and HEK293T cell lines were obtained from the American Type Culture Collection. All cells were maintained at 37 °C with 5% CO₂ in high-glucose DMEM medium supplemented with L-glutamine, sodium pyruvate (Gibco, 11995065), and 10% FBS (Sigma-Aldrich, F4135).

### Cell strain construction

The A375-dCas9-BFP-KRAB cell strain was constructed by transducing A375 cells with lentivirus produced using pHR-SFFV-dCas9-BFP (Addgene, 46910) at a multiplicity of infection (MOI) of 0.2. BFP+ cells with the top 15% fluorescence intensity were sorted for expansion. Enrichment sorting was performed regularly to maintain the Cas9 expression level in the cell strain.

### Plasmid and Lentivirus Construction

Single sgRNA expression construct cloning and lentivirus packaging were performed similarly to our previous work(*8*). In brief, the CROPseqOPTI vector with an additional EGFP marker (*CROPseqOPTI-GFP*) was digested with BsmBI (NEB, ER0451) to remove the filler sequence between the U6 promoter and the sgRNA scaffold. Pre-annealed phosphorylated double-stranded DNA with an sgRNA sequence and complementary sticky ends was ligated into the digested backbone. Lentivirus was packaged by co-transfecting the sgRNA-expressing plasmid, psPAX2 plasmid, and pMD2.G plasmid into HEK293T cells at high confluency. After washing the cells and changing the medium, virus-containing supernatants were harvested 24 hours later. The functional titer of the viruses was measured by assessing the proportion of fluorescent marker-positive cells after transducing target cell lines with varying amounts of lentivirus.

### Gene selection and sgRNA evaluation

To identify potential gene candidates whose loss of function might confer resistance to Vemurafenib treatment in melanoma cells, we analyzed previously published bulk CRISPR screens with similar experimental settings(*25–28*). From these datasets, we identified 183 genes with an FDR < 0.1 in at least one screen. Candidate genes for knockdown were further filtered based on their endogenous expression levels; 143 genes with expression levels ≥ 1 log2 counts per million (CPM) in our in-house A375 EasySci-RNA data were retained.

We referred to the sgRNA library from Genome-wide Perturb-seq, which also utilized a paired sgRNA expression system but employed a different chemistry. We further examined the sgRNAs and retained only those meeting the following criteria: (1) unique STAR mapping to the reference genome (GRCh38.p13, GENCODE), and (2) knockdown fold-change < 0.5 (i.e., the fold-change of target gene expression between cells with a target sgRNA and cells without a target sgRNA), as reflected in the Genome-wide Perturb-seq gene expression data. For target genes not found in the Genome-wide Perturb-seq or in cases where sgRNA inefficiency was observed in the original dataset, we used newly designed paired sgRNA libraries from (*85*).

### CRISPR screen experimental procedure

For each replicate, 5 million A375-dCas9-BFP-KRAB cells were seeded into a 10 cm dish. The next day, the drug-resistance lentivirus library was added to the dish at an MOI of 0.05 with 8 ng/ml Polybrene (Sigma-Aldrich, TR-1003). GFP+ BFP+ cells were sorted and reseeded into a 6 cm dish for expansion. After 48 hours, the medium was replaced with fresh DMEM containing either 0.1% DMSO (VWR, IC0219605525) or 1 μM Vemurafenib in DMSO (Selleck, S1267) to initiate the screen (day 0). The culture medium was refreshed every other day.

On day 6, after 5-EU labeling, cells were trypsinized and collected for single-cell library preparation. To minimize the impact of experimental handling on nascent transcriptomes, cells were processed on ice immediately after collection and fixed promptly. Detailed step-by-step *PerturbFate* protocols are provided in the Supplementary Materials.

For the bulk screen, cells were trypsinized and collected on days 6 and 14. Cell pellets were stored at −20 °C until further processing. Genomic DNA was extracted from each sample (Zymo, D3024) and split equally into multiple PCR reactions, with a maximum of 1 mg of DNA per reaction for sgRNA expression cassette amplification. After 20 cycles of amplification, the PCR products were purified (Zymo, D4014), and the target band was extracted from an agarose gel (Zymo, D4007) for amplicon sequencing.

### Sequencing data processing

For the bulk CRISPR screens, BCL files were demultiplexed into FASTQ files based on index 7 barcodes. Reads from each sample were further extracted using index 5 barcodes. Each read pair was matched against two constant sequences (Read1: 11–25 nt; Read2: 11–25 nt) to retain reads with the correct library structure, allowing for a maximum of one mismatch. sgRNA sequences were then extracted from filtered read pairs (sgRNA sequence 1: 25–44 nt of Read1; sgRNA sequence 2: 84–103 nt of Read2). sgRNA identities were assigned based on the reference sgRNA sequences, permitting no mismatches. sgRNA count matrices were constructed using Python 3.8.

For PerturbFate (whole transcriptome + chromatin accessibility + sgRNA), BCL files were demultiplexed into FASTQ files using P7 PCR barcodes. Each FASTQ file contained all RNA and ATAC reads from the same indexing PCR well. During barcode extraction, reads from the oligo-dT/randomN RNA and ATAC modalities were split into separate files based on modality-specific adapter sequence matching. Read1 matching the “TG” sequence at the 41–42 nt position was classified as transcriptome data. Oligo-dT-originated RNA reads required at least 10 consecutive T bases at positions 51–60 nt, while randomN-originated RNA reads did not exhibit this pattern. ATAC read pairs were identified by the Tn5 ME sequence at the 41–50 nt position of Read1.

For the RNA reads, cell barcodes and UMI sequences from Read1 were appended to the headers of each Read2 FASTQ file, and only Read2 was used for subsequent steps. Poly(A) sequences, adapter sequences, and low-quality bases were trimmed from Read2 of oligo-dT reads, while 14 additional bases (potential priming hexamer and UMI sequences) were removed from randomN reads using Trim_Galore (0.6.7). After this step, oligo-dT and randomN reads were combined for further processing. STAR (2.7.9a) was used to map reads to the hg38 reference genome (GRCh38.p13, GENCODE). Reads with a mapping score ≥30 were selected using SAMtools (1.13). Single-cell deduplication was performed based on UMI sequences and alignment locations, and retained reads were split into SAM files per cell. Finally, single-cell gene-to-cell raw count matrices were constructed by assigning reads to genes.

For the ATAC reads, cell barcodes from RNA Read1 were appended to the headers of both ATAC Read1 and Read2 FASTQ files. Adapter sequences were removed in paired-end mode using Trim_Galore (0.6.7). STAR (2.7.9a) was used to map read pairs to the hg38 reference genome (GRCh38.p13, GENCODE). Picard (2.27.4) was used to remove PCR duplicates, and TSV files containing genomic locations and full cell barcodes were generated for SnapATAC2 (2.6.1) (*56*).

sgRNA reads were matched against constant sequences in Read1 and Read2, allowing a maximum of one mismatch. For each filtered read pair, the cell barcode, sgRNA sequence, and UMI were extracted from designed positions. Extracted sgRNA sequences with at most one mismatch from the sgRNA library were corrected. UMI sequences were used for deduplication by collapsing identical UMIs for each individual sgRNA under a unique cell barcode. Cells with sgRNA UMI counts exceeding 5 were retained, and the sgRNA × cell count matrix was constructed.

For PerturbFate (nascent transcriptome + pre-existing transcriptome + chromatin accessibility + sgRNA), most computational steps were the same as described above, with minor modifications. In this protocol, nascent and pre-existing transcriptomes from the same cell were amplified separately using distinct P7 PCR barcodes and processed as separate cells (two groups of reads with distinct cell barcodes). These were matched after single-cell gene-to-cell raw count matrices were generated. Additionally, because ATAC DNA underwent secondary tagmentation, the pipeline was modified to process single-end Read1 of ATAC fragments for chromatin accessibility quantification. The sgRNA library from the same cell was amplified from both nascent and pre-existing transcriptome PCR wells but shared a unique cell barcode for matching, achieved by adding the same barcoded sgRNA inner P7-U6 primer to paired nascent and pre-existing transcriptome PCR wells.

ATAC peaks were called for each treatment condition with a q-value cutoff of 0.01 using MACS3 (3.0.0b3) (*86*) embedded in SnapATAC2 (2.6.1). Peaks were resized to 500 bp centered at the summit and iteratively merged into a union peak set, resulting in 758,365 peaks in the final set. A peak-to-cell raw count matrix was generated using SnapATAC2 and converted into a sparse matrix in R for downstream analyses.

### Calculation of Growth Phenotypes in Bulk CRISPR Screens

Raw sgRNA counts from the plasmid library, as well as from cells in the DMSO and Vem conditions, were normalized by the total counts to account for sequencing depth. Perturbations relative to NTC counts were calculated by dividing the depth-normalized sgRNA counts by the sum of depth-normalized NTC counts. Relative counts from replicates were then averaged for growth phenotype calculations. To assess whether perturbed cells outgrew NTC cells under the Vem condition, fold changes in relative counts between the Vem library and the plasmid library were calculated and log2-transformed (*i.e.,* Vem log2(sgRNA enrichment)). The Vem log2(sgRNA enrichment) value indicates whether perturbed cells exhibit a growth advantage over NTC cells in response to Vem treatment. To further compare changes in cell growth under Vem treatment, the ratio of fold changes in relative counts between the Vem library and the DMSO library was calculated and log2-transformed (*i.e.,* Vem - DMSO log2(sgRNA enrichment)). The Vem - DMSO log2(sgRNA enrichment) value reflects whether perturbed cells gained a greater growth advantage in the Vem condition compared to the DMSO condition.

### sgRNA singlets identification and knockdown efficiency examination

sgRNA singlets were identified using two additional cutoffs: a Max-to-Total proportion ≥ 50% and a Sec-to-Max ratio ≤ 0.3. The Max-to-Total proportion represents the percentage of UMI counts for the most abundant sgRNA relative to the total sgRNA UMI counts from a single cell. The Sec-to-Max ratio represents the ratio of UMI counts for the second most abundant sgRNA to the UMI counts for the dominant sgRNA in a single cell. We benchmarked our method against a recent Python package that implements multiple statistical inference methods for single-cell sgRNA identities (Braunger and Velten, 2024) and observed extremely high consistency. We retained only knockdown populations with more than 50 cells in each treatment condition and those in which the pseudobulk expression level of the target gene was less than 60% of that observed in NTC cells.

### Differential gene expression (DEGs) and differential accessible peaks (DAPs) analyses

Whole-transcriptome differential expression analysis was performed similarly to our previous work (*3*, *8*, *87*). For each knockdown-to-NTC DEG test, cells with fewer than 100 detected genes and features present in fewer than 25 cells were filtered out. The analysis was conducted using Monocle2 (2.34.0) (*88*). The differentialGeneTest() function was implemented with the number of genes detected per cell as a covariate. The fold change was calculated by dividing the average normalized single-cell gene expression CPM in knockdown cells by that in NTC cells. Differentially accessible peaks were identified using the diff_test() function in snapATAC2 with default parameters.

### UMAP embedding on pseudobulk perturbations

Pseudobulk knockdown population whole-transcriptome matrices and ATAC peak count matrices were constructed by aggregating all cells with the same sgRNA identity within each treatment condition. Seurat (4.2.0) (*89*) was used for normalization, scaling, PCA dimensionality reduction, and UMAP visualization. Single-cell level MAST differential expression analysis of knockdown cells versus NTC cells was performed to obtain gene features for dimensionality reduction. Perturbations with at least one significant DEG were retained, and the top 20 DEGs with the highest absolute fold changes from all tests were combined. The top 15 principal components were used for UMAP. For ATAC pseudobulk dimensionality reduction, 300,000 highly variable peaks were selected, as implemented in snapATAC2. The top 30 principal components were used as input for UMAP.

### ATAC peaks - nascent gene expression linkage identification

The FigR (1.0.1) package (*38*) in R was used for the analysis. To account for the potential effects of genetic and chemical perturbations on global transcriptional activity, we employed a specialized normalization strategy to adjust for sequencing depth. While NTC cells in the DMSO condition were normalized by counts per 10k, other perturbed cells were normalized using a scaling factor of 10k x (mean UMIs in this population/mean UMIs in NTC cells in the DMSO condition). Given that gene regulation patterns of non-coding genes remain largely understudied, we focused exclusively on protein-coding genes. Only protein-coding genes annotated in the hg38 GTF file (GRCh38.p13, GENCODE) were retained. The ATAC peak-to-cell raw count matrix was filtered to include only peaks detected in at least 10 cells. The log-transformed, normalized nascent gene expression matrix and the filtered ATAC peak-to-cell raw count matrix were used as input for the association test. Gene-peak linkages with p-values (pvalZ) ≤ 0.05, as specified in the FigR manual, were retained as significant linkages.

### Cell trajectory and nascent transcriptome-based RNA velocity inference

Since whole transcriptomes of single cells remain the most robust single-cell modality, they were used for diffusion map construction and cell order (pseudotime) inference using the destiny package (3.20.0) in R. To highlight the impact of genetic and chemical perturbations on the melanoma phenotype switch, well-established gene sets defining multiple melanoma phenotypic states (*35*) were used as features. Genes expressed in at least 100 cells in each condition were retained for dimensionality reduction. The top 20 principal components were used to construct the diffusion map with the DiffusionMap() function, and cell order was subsequently calculated using the DPT() function. Phenotype gene set scores in single cells were calculated by averaging the scaled expression values of constituent genes, and cells were annotated according to the highest gene program score in each bin along the trajectory. To improve clarity and ensure the statistical robustness of downstream analyses, transient plastic phenotype labels were excluded when annotating cells in the DMSO condition. Instead, cells were categorized as one of “Melanocytic,” “Neural crest-like,” or “Undifferentiated.” For Vem-treated cells, no phenotype labels were assigned; instead, we focused on their transitions along the trajectory, reasoning that their transcriptomic states underwent complex and subtle changes after treatment, making them unsuitable for categorization into untreated phenotypes.

To integrate single-cell nascent transcriptomes into the diffusion map, we assumed that cell state transitions were not completed within one hour. This assumption aligns with the observation that RNA half-lives in human cells are typically much longer than one hour(*90*). Whole-transcriptome gene count matrices of cells were concatenated with pre-existing transcriptome gene count matrices of cells (which had matched single-cell nascent transcriptomes), and a joint diffusion map was constructed. Manual curation revealed that DC2 and DC3 captured the phenotypic state of cells across both chemistries. We constructed a k-nearest neighbor model using DC2 and DC3 values with the FNN package (1.1.4.1) in R. The inferred cell order value for each pre-existing transcriptome cell was assigned by averaging the real cell order values of its three nearest neighbor whole-transcriptome cells. This approach demonstrated highly consistent sgRNA abundance distributions (**Fig. S6A**).

The scvelo (0.3.2) package(*91*) in Python was used to infer RNA velocity from nascent transcriptomes. To match nascent transcriptomes to whole-transcriptome cells, we performed integration as described above. Each whole-transcriptome cell was matched to a nascent transcriptome by identifying the two nearest pre-existing transcriptome neighbors and averaging their log-transformed normalized nascent transcriptomes. Normalized whole and nascent transcriptome gene expression matrices, diffusion map coordinates of whole-transcriptome cells, and principal components computed with destiny were imported into the adata object. Instead of using the unspliced-to-spliced ratio, we used the new-to-total ratio of genes (*92*) to compute velocities. The top 3,000 genes were selected, and the top 20 principal components were used to compute moments and connectivity. Velocities were generated in stochastic mode.

### Significance examination of perturbed cells in the trajectory

The ks.test() function in R was used to identify perturbed cells that exhibited significant shifts along the trajectory compared to NTC cells in both conditions. Cell order values for each perturbation were compared to those of NTC cells in the corresponding treatment condition, and differences in mean cell order values between perturbations and NTC cells were used to determine the direction and extent of the shift. The computed two-sided p-values were adjusted using FDR correction. For most analyses of Vem resistance, we retained only the perturbations in the Vem condition that displayed consistent growth and molecular phenotypes (*i.e.,* shift trends along the trajectory).

To compare the relative transition distance between NTC and other perturbations upon Vemurafenib treatment, the wilcox.test() function in R was used in a similar manner. For both DMSO and Vemurafenib treatment conditions, single-cell order values of perturbed cells were adjusted relative to the averaged cell order value of the NTC cell population, and the difference between these two distributions was examined using the wilcox.test() function followed by FDR correction.

### Identification of DEGs, DA motifs, and DAPs along the cell trajectory

The TradeSeq (1.20.0) package (*93*) in R was used to infer statistically significant DEGs, DA motifs, and DAPs along the melanoma phenotype switch trajectory. For whole-transcriptome and nascent-transcriptome DEG identification, raw gene count matrices (whole transcriptome and nascent transcriptome) and cell order values inferred by destiny (for whole transcriptome cells) or by the integration mentioned above (for nascent transcriptome cells) were used as input to the fitGAM() function. Lineage weights for all cells were set to 1, as only a single trajectory was analyzed. The statistical significance of associations between expression levels and cell order was calculated using the associationTest() function. Each trajectory was divided into 40 bins based on cell order values. Genes with an FDR < 0.01 in both whole-transcriptome and nascent-transcriptome analyses were considered for gene regulatory network (GRN) construction. Genes with an FDR < 0.01 and an absolute log2 fold change > 1 were selected for heatmap visualization in **Figure 2F**. Motif counts in single cells were obtained using the motifmatchr package (1.28.0) in R. The peak-to-cell raw matrix was used as input, sequences of peaks were scanned, and motifs from the JASPAR database (*94*) were matched against sequences in open peaks of cells. Significant motifs (FDR < 0.01) were included in GRN construction and heatmap visualization. For DAP analysis along the drug-resistance trajectory, data sparsity in single-cell ATAC and program memory requirements were addressed by binning the trajectory as follows: 1,000 cells on the leftmost (melanocytic) side of the trajectory were divided into 10 bins, 300 cells on the rightmost (undifferentiated) side were divided into 3 bins, and the remaining cells in the middle were divided into 10 bins. For bins in the middle containing more than 100 cells, 100 cells were randomly sampled. ATAC raw peak counts were aggregated for each bin, and only peaks with non-zero counts in more than one-third of the bins were retained. Subsequent steps followed the methodology described above. Motif enrichment in DAPs was performed using HOMER (5.1) (*95*).

### Melanoma phenotype switch GRN inference

The GRN inference was conducted through the following steps. First, we determined the initial and terminal states of each phenotype switch, which defined the directionality of the temporal constraints and the cell order window to use. Cells in the DMSO condition initially exhibited a Neural crest-like phenotype and transdifferentiated into either melanocytic (DMSO Neural crest-like to Melanocytic trajectory) or undifferentiated phenotypes (DMSO Neural crest-like to Undifferentiated-1 trajectory and DMSO Undifferentiated-1 to Undifferentiated-2 trajectory). The bin containing the median cell order value of DMSO NTC cells was set as the root for both the DMSO Neural crest-like to Melanocytic trajectory and the DMSO Neural crest-like to Undifferentiated-1 trajectory. Additionally, the bin where the threshold value between gene clusters 7 and 8 (**Figure 2F**) was reached was set as the root for the DMSO Undifferentiated-1 to Undifferentiated-2 trajectory. For cells under Vemurafenib treatment, we observed a strong trend of transdifferentiation toward the melanocytic side of the trajectory, particularly from the normal WT Neural crest-like phenotype to the inactivated Neural crest-like phenotype (Vem-sensitive trajectory). Conversely, drug-resistant cells transitioned from the inactivated Neural crest-like phenotype to the undifferentiated phenotype (Vem-resistance trajectory). The root of the Vem-sensitive trajectory was set at the bin containing the median cell order value of DMSO NTC cells, while the root of the Vem-resistant trajectory was set at the bin immediately following the one containing the median cell order value of Vem NTC cells.

We performed K-means clustering (k = 10) on all significant DE steady-state and nascent z-scored gene expressions along the trajectory. Genes were filtered by correlating steady-state gene expression with nascent gene expression at the cluster level along the trajectory. Steady-state and nascent gene clusters with correlations ≥ 0.6 were matched, and genes present in both counterparts of matched clusters, with nascent gene expression peaking either before or simultaneously with steady-state gene expression in the direction of transdifferentiation, were retained as GRN target gene candidates. We then determined the association between TF motifs and genes. If TF motifs identified by motifmatchr were found in peaks significantly associated with the nascent expression of a gene, a TF-gene linkage was considered. Significant z-scored TF motif counts along the trajectory were clustered using K-means (k = 10), and only TF motif counts that showed consistent trends with their steady-state gene expression (correlation ≥ 0.4 and manual curation) were retained. Motif clusters and nascent gene expression clusters were matched in the same manner as described above. TF-gene linkages were retained only if they met the following criteria: (1) the associated peaks of a gene contained at least one motif of the TF from its matched counterpart motif cluster, and (2) the peak of motif opening along the trajectory occurred before or at the same time as the peak of nascent gene expression.

For each transdifferentiation trajectory segment (*e.g.,* DMSO Neural crest-like to Undifferentiated-1 trajectory, DMSO Undifferentiated-1 to Undifferentiated-2 trajectory, DMSO Neural crest-like to Melanocytic trajectory, Vem-sensitive trajectory, and Vem-resistant trajectory), only clusters from all modalities (*i.e.,* steady-state, nascent transcriptome, and motif) that exhibit peak values within the segment were considered for segment-specific GRN construction.

### Multi-modal regulon scoring analysis

A regulon is defined as a group of target genes co-regulated by an upstream TF. We propose an integrated score (Regulon Score) to quantify the overall transcriptional activity of this gene group in a group of cells that received the same sgRNA:

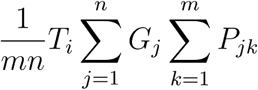

Where *T* ɛ (*T*_1_, *T*_2_, … *T_l_*) is the log-transformed, normalized pseudobulk steady-state expression level of the regulator TF *l* in a perturbed population; *G* ɛ (*G*_1_, *G*_2_, … *G_n_*) is the log-transformed, normalized pseudobulk nascent expression level of a target gene regulated by the TF in a perturbed population; *P_j_* ɛ (*P_j_*_1_, *P_j_*_2_, … *P_jm_*) is the log-transformed, normalized ATAC-seq counts at a peak associated with the target gene *j*, which contains the motif of TF *i* in a perturbed population.

To identify the pattern of inter-regulon co-regulation in Vem resistance, we selected perturbations in the Vem condition that exhibited a significant shift toward the undifferentiated end of the trajectory with consistent growth phenotypes, as well as perturbations in the Vem condition that showed no significant shift in the trajectory. The fold change in Regulon Scores between perturbations and NTC was calculated, and Pearson correlations of these fold changes between regulons across the selected perturbations were computed. Distances between regulons were calculated, and hierarchical clustering was performed on the resulting correlation coefficient matrix. Clusters of regulons were then defined using the cutree() function.

To assess the statistical significance of the Regulon Score fold changes for each perturbation compared to NTC in the Vem condition, we implemented a permutation strategy. For each regulon in each perturbation, the order of *G* was randomly permuted 500 times, and the permuted Regulon Scores were divided by the real Regulon Score of the NTC. This process generated a fold change background distribution for each regulon in each perturbation. Two-sided empirical p-values were calculated for each regulon in each perturbation and subjected to FDR correction. Regulon Scores with FDR < 0.1 were considered significant.

### Cell order - gene signature correlation analysis and YAP ChIP-seq analysis

To identify potential contributors to the cell state shift in the trajectory, oncogenic signatures from MSigDB (*96*) in Vem-treated single cells were scored using the AddModuleScore() function in Seurat. Pearson correlations between cell order values and gene signature scores were calculated. Gene signatures with significant correlations (FDR < 0.01) were further tested with 1,000 permutations, followed by FDR correction, to confirm that the observed correlations were not due to random chance. YAP indirect binding peaks were obtained from a previous report (*65*). The PScan-ChIP tool (1.3) (*97*) was used to analyze local motif enrichment within these peaks, identifying potential direct downstream signal transmitters of YAP.

### TF motif deviation scoring analysis

The ChromVAR (1.28.0) package(*98*) in R was used to compute TF motif deviation scores for individual cells. Motif scores for each cell were calculated using peaks that were open in at least 10 cells. GC-matched backgrounds were then added using the addGCBias() function, and the computeDeviations() function was executed to obtain motif deviation scores. For each treatment condition, the deviation score distribution of each motif was compared to that of NTC cells. Simultaneously, the steady-state DEG results between perturbed populations and NTC cells were intersected with the motif score tests. Only TFs that were significant in the DEG analysis and exhibited a minimum level of gene expression changes were retained. Two sided p-values for these motifs were further corrected for false discovery rate (FDR). Significant motifs were defined as those with an FDR < 0.1, with enrichment or depletion determined based on the median z-score differences between perturbed cells and NTC cells.

### Transcriptome and TF deviation score-based clustering

To cluster perturbed populations, the top 200 whole-transcriptome DEGs with the highest fold changes were extracted and combined into a unified gene set. Pairwise Pearson correlation coefficients between perturbations were calculated based on the fold changes of these populations compared to NTC cells. Perturbation clustering based on motifs was performed using the median deviation score differences between the perturbed populations and NTC cells. Deviation score differences were set to 0 if a motif was not present in the original analysis output.

### Mediator Complex perturbations associated gene program analyses

We established the following criteria to identify gene signatures in Oncogenic Signatures that might account for heterogeneous phenotypic responses to genetic perturbations in the Mediator Complex and Vem treatment: (1) Under the DMSO condition, compared to NTC cells, the scores of gene signatures in both MED15 and MED24 showed no significant changes (based on Wilcoxon tests with FDR corrections); (2) Under the Vem condition, compared to NTC cells, the scores of gene signatures in both MED15 and MED24 showed significant changes (based on Wilcoxon tests with FDR corrections).

### Published paired RNA-seq and SOX10 ChIP-seq analyses

Genomic SOX10 binding profiles and paired bulk gene expression were analyzed in a previous study(*76*) using another melanoma cell line harboring the *BRAF^V600E^* mutation, the 501Mel cell line, treated with a different MAPK signaling inhibitor, AZD6244. SOX10 ChIP-seq and bulk RNA-seq FASTQ files were downloaded and reprocessed from NCBI BioProject PRJNA515302. For bulk RNA-seq processing, adapter sequences were removed in paired-end mode using Trim_Galore (0.6.7). Reads were mapped to the hg38 reference genome (GRCh38.p13, GENCODE) using STAR (2.7.9a). Reads with mapping scores <10 were filtered out using SAMtools (1.13). PCR duplicates were removed with Picard (2.27.4), and gene counts were generated using featureCounts (2.0.1). For ChIP-seq processing, adapter sequences in FASTQ files were trimmed using cutadapt (3.4). Reads were aligned to the hg38 reference genome using Bowtie2. PCR duplicates were removed using Picard (2.27.4), and peaks were called using MACS2 (2.2.9.1) with a minimum q-value threshold of 0.01.

ChIP-seq peaks called from replicates were combined and resized to 500 bp, extending from the center. A total of 2,003 and 7,159 peaks were identified for the DMSO and AZD6244 conditions, respectively, with 1,878 overlapping peaks. Significantly upregulated DEGs with an FDR < 0.05 were identified using the DESeq2 package (*99*) in R. To identify potent MAPK/ERK inhibitor-responsive SOX10 targets, the nearest genes within 10 kb of 5,281 AZD6244-specific SOX10 binding peaks were identified using the closest command from bedtools (2.30.0) and intersected with the significantly upregulated DEGs. Genes from this intersection that also met the following criteria were considered SOX10 direct targets responsive to Vem treatment: (1) the maximum steady-state expression CPM in either DMSO NTC cells or Vem NTC cells was ≥ 20, (2) the gene was significantly upregulated in NTC cells upon Vem treatment, and (3) the fold change was ≥ 1.25. Ratios of these genes between the Vem-to-DMSO fold change in perturbed cells and the Vem-to-DMSO fold change in NTC cells were calculated to assess the strength of SOX10-mediated gene activation upon Vem treatment.

### Hybridization Chain Reaction (HCR) staining on intracellular RNA

The HCR reaction was performed to quantify intracellular AXL expression levels. Probe pairs targeting AXL mRNA were designed in-house, and the following steps were carried out based on a published protocol(*100*). In brief, cells were fixed in 4% formaldehyde at room temperature for 1 hour, followed by ethanol permeabilization at −20°C overnight. Hybridization was performed at 37°C overnight using 40 nM per probe. Hairpins conjugated with an Alexa647 fluorophore (Molecular Instruments, HCR Gold Amplifier) were used for signal amplification at room temperature overnight. Flow cytometry analysis was performed to record intracellular signals.

## Acknowledgments

We thank the members of the Cao lab, Artem Khan from Rockefeller University, and Erting Tang from the University of Chicago for their helpful discussions and feedback. We also thank the members of the Rockefeller University High-Performance Computing Core, Genomics Resource Center, and Bioinformatics Resource Center for their support. Diagrams were created with BioRender.com.

## Author contributions

J.C. and Z.X. conceived the study. Z.X. developed and optimized the *PerturbFate* protocols. Z.X. performed all experiments and computational analyses with insights from Z.L., A.U., and A.A.. Z.X. and A.L. conducted the HCR experiment. J.Z. provided insights into structural biology and replotted the protein structure. J.C., W.Z., and Z.X. wrote the manuscript with input from all co-authors.

## Data availability

MED1 and YAP ChIP-seq data were downloaded from the GEO dataset GSE62275. MED1 and MED12 ChIP-seq data under *AAVS1-* or *MED12-*KO conditions were downloaded from the GEO dataset GSE174282. Single-cell RNA-seq data from NTC and *SOX10*-KD melanoma patient-derived cell cultures were downloaded from the GEO datasets GSM3946507, GSM3946510, GSM3946511, GSM3946512, and GSM3946513. Bulk ATAC-seq data from NTC and *SOX10*-KD melanoma patient-derived cell cultures were downloaded from the UCSC Genome Browser track hub via the following link: http://ucsctracks.aertslab.org/papers/wouters_human_melanoma/hub.txt. Genomic SOX10 ChIP-seq data and paired bulk RNA-seq data from the Mel501 cell line, with or without AZD6244 treatment, were downloaded from Bioproject PRJNA515302.

## Supplementary information

### Supplementary Text1

The detailed cost estimation of *PerturbFate* single-cell sequencing library preparation.

---

**Fig. S1:**
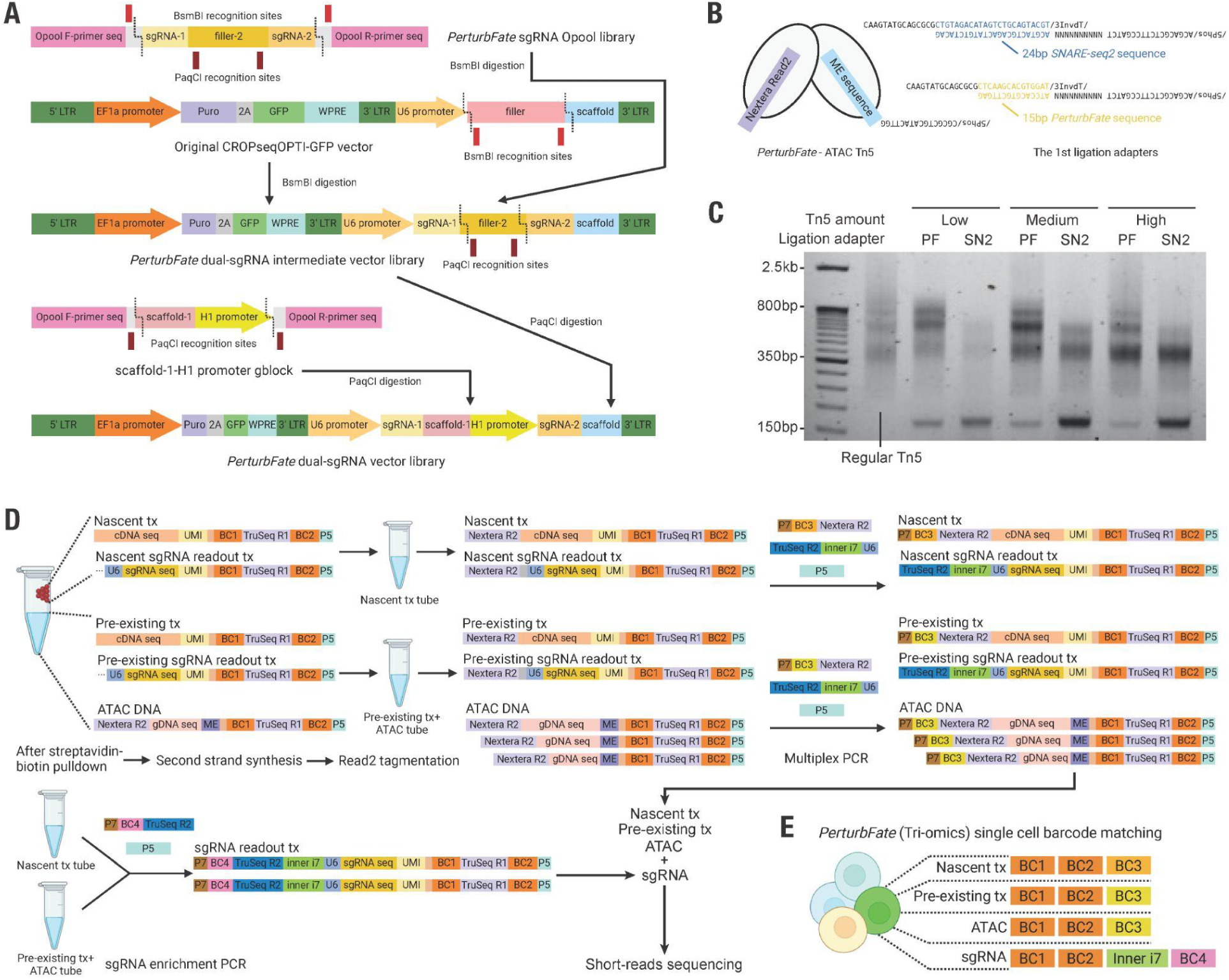
Schematics of *PerturbFate* Vector Library Construction and Sequencing Library Preparation. **(A)** Schematics of *PerturbFate* vector library construction. After the sgRNA oligo pool is amplified into double-stranded DNA, BsmBI digestion is performed on both the amplified library and the backbone. sgRNA pairs separated by a filler sequence, are cloned into the *CROP-seq-OPTI* backbone by ligation. The filler sequence is subsequently replaced by the sgRNA scaffold and the hH1 promoter through another round of PacCI digestion and ligation. **(B)** Schematics of the *PerturbFate*-ATAC Tn5 oligo structure. The two ends of Tn5 consist of the Nextera Read 2 sequence and a ligation adapter flanking the double-stranded Mosaic End (ME) sequence. SNARE-seq2 ligation sequence(*18*) and in-house ligation sequence were tested. **(C)** Representative gel image of ATAC libraries performed using Tn5 with different adapters and different amounts. 100,000 cells underwent tagmentation using 2.5/5/10uL of regular Tn5 with Nextera Read1/2 sequences, Tn5 with Nextera Read2 and *PerturbFate* ligation adapter, or Tn5 with Nextera Read2 and *SNARE-seq2* (SN2) ligation adapter. After the ligation of ATAC libraries prepared with *PerturbFate* and *SNARE-seq2* Tn5, all libraries were amplified and visualized. **(D)** Schematics of *PerturbFate* sequencing library structures during library preparation. After cell lysis and biotin-streptavidin pulldown, the supernatant containing ATAC fragments and pre-existing transcriptomes, and beads carrying nascent transcriptomes, are separated into two tubes. Second-strand synthesis and cDNA Read2 tagmentation of both compartments are performed separately. Multiplex PCR then adds matched third cell barcodes to the nascent/pre-existing transcriptome and ATAC libraries while incorporating a unified inner i7 barcode into the sgRNA library in both tubes. A small portion of the PCR product from the matched PCR tubes is used for another sgRNA enrichment PCR, which adds fourth cell barcodes to the sgRNA library. **(E)** Schematics of *PerturbFate* Tri-omics post-sequencing cell barcode matching. For each single cell with multiple molecular modalities, all modalities share the first two cell barcodes. The ATAC and pre-existing transcriptome share the same cell barcode 3, while the nascent transcriptome has a matched but distinct cell barcode 3, as it is amplified in a separate PCR tube. For the sgRNA modality, each inner i7-barcode 4 combination corresponds to a unique barcode 3 pair from other modalities.

**Fig. S2:**
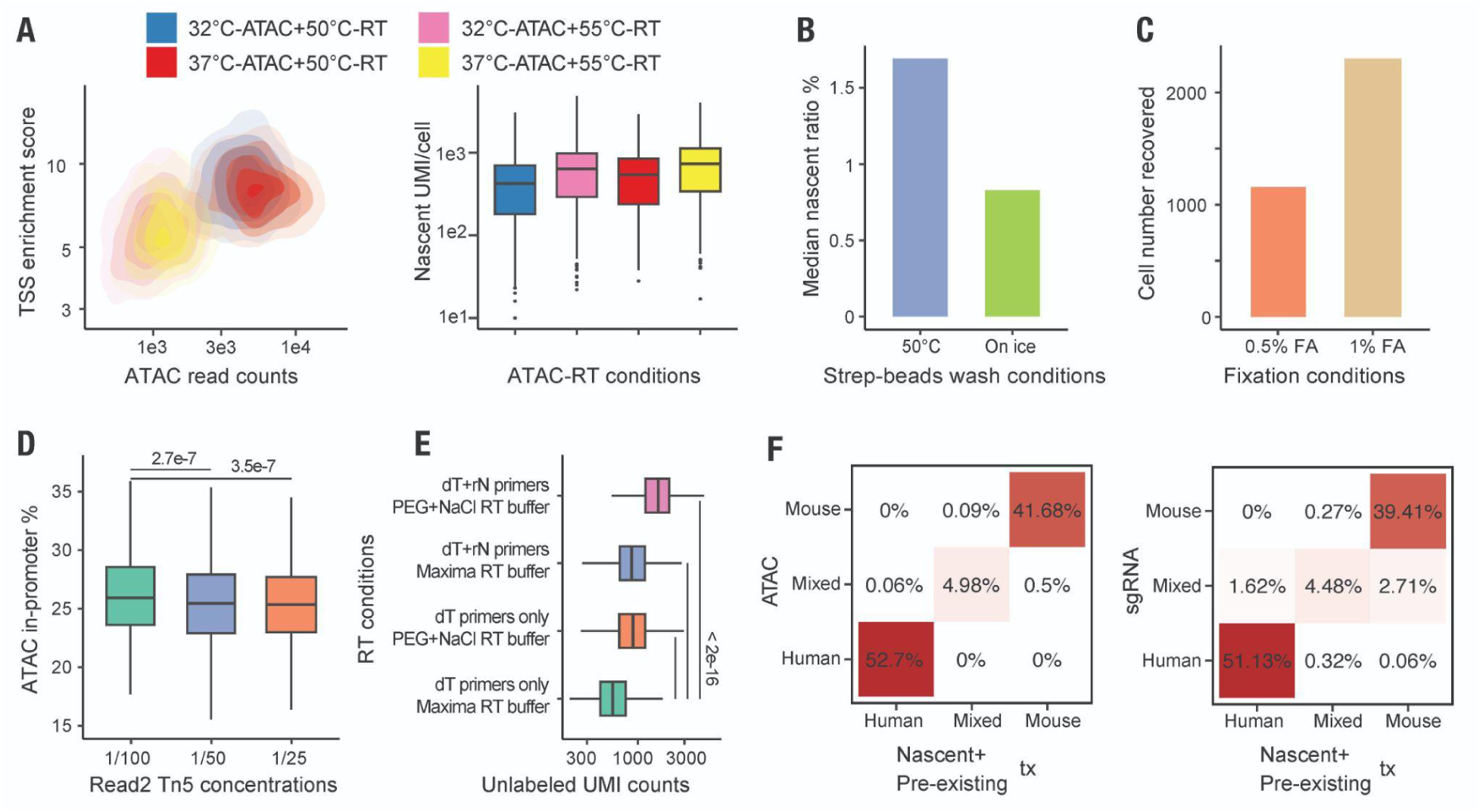
Comprehensive optimizations substantially improved the quality of *PerturbFate*. **(A)** Density plot showing the number of reads and TSS enrichment of single-cell ATAC libraries recovered from different reaction temperatures (left panel) and box plot showing the number of UMIs in single-cell nascent transcriptomes recovered from different reaction temperatures (right panel). In this experiment, tagmentation in bulk was performed at either 32°C or 37°C to assess potential RNA degradation during the ATAC reaction. Additionally, the maximum reaction temperature for reverse transcription was set to either the default 55°C or reduced to 50°C to evaluate its effect on ATAC fragment loss. No observable RNA degradation was detected at 37°C compared to 32°C; however, ATAC quality was severely compromised when reverse transcription was performed at 55°C, potentially due to substantial DNA leakage. **(B)** Bar plot showing the mean percentage of nonspecific nascent reads pulled down from unlabeled cells under different streptavidin bead pulldown conditions. To minimize nonspecific contamination during the biotin-streptavidin bead pulldown, streptavidin beads were washed either on ice or at 50°C following scEU-seq(*19*). No additional benefit was observed from increasing the washing temperature. **(C)** Bar plot showing the number of multi-modal cells recovered from the experiment with cells fixed under different conditions. A mixture of cells fixed by 0.5% or 1% Formaldehyde was used for the experiment. **(D)** Box plot showing the effect of Read2 Tn5 concentration on ATAC library quality. To account for potential Read 2 tagmentation on ATAC fragments during the cDNA tagmentation step, only Read1 of the ATAC library was used to infer chromatin accessibility. Increasing the Tn5 concentration slightly decreased single-end ATAC quality but remained overall stable. **(E)** Box plot showing UMI of single-cell pre-existing transcriptomes recovered from different reverse transcription conditions. Incorporating random hexamer primers and replacing the original Maxima reverse transcription buffer with the optimized buffer from SMART-seq3(*20*) significantly improved the transcriptomic UMI. **(F)** Heatmaps showing the consistency of species assignment for single cells at the multi-modal level. A mixture of human HEK293T cells and mouse NIH/3T3 cells, each transduced with a different sgRNA, was used for profiling. RNA and ATAC modalities were aligned to a human-mouse hybrid genome for species assignment, while sgRNA reads were demultiplexed to assess the single-cell purity of the library preparation. Boxes in box plots indicate the median and interquartile range (IQR), with whiskers indicating 1.5× IQR.

**Fig. S3:**
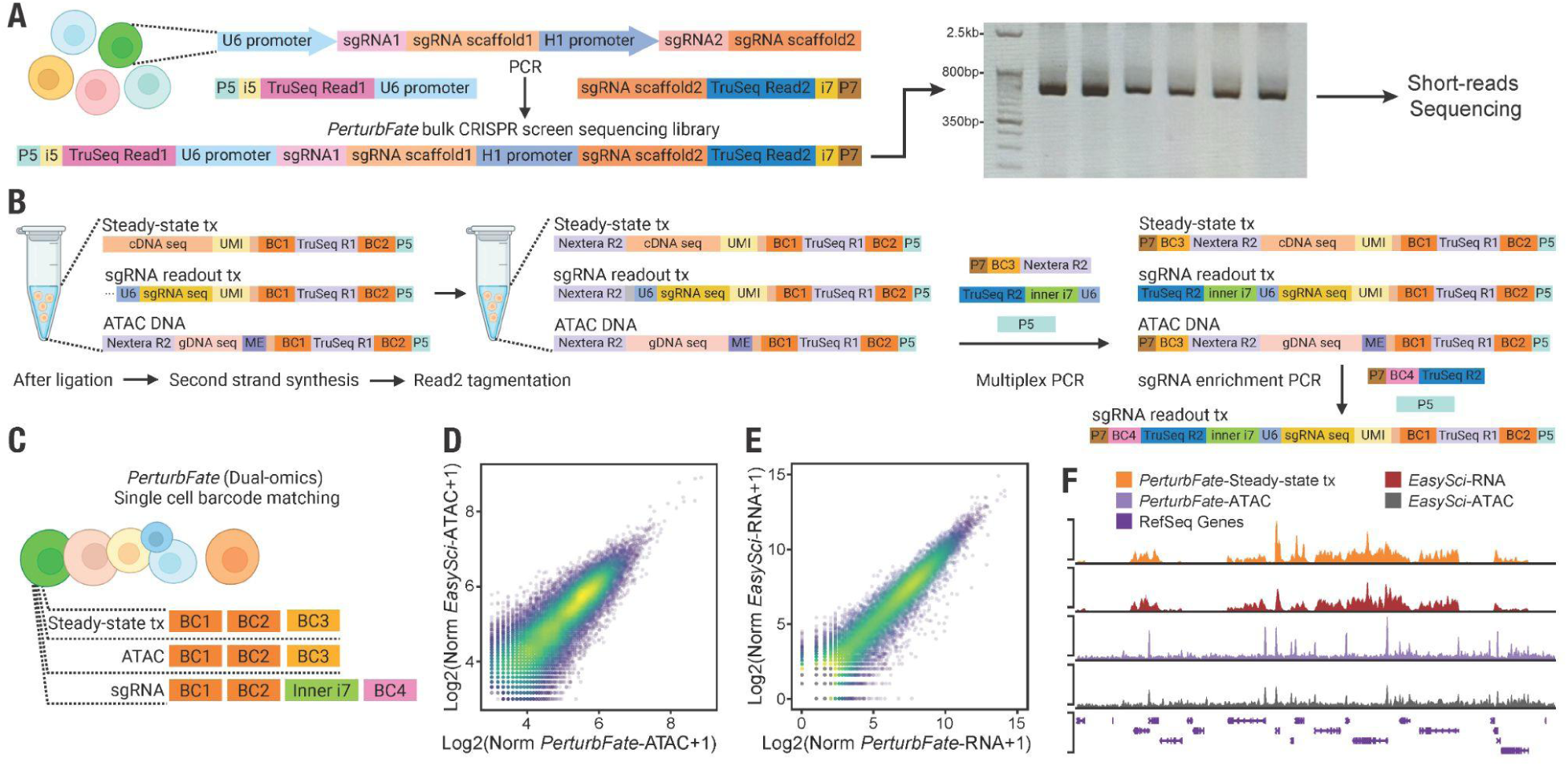
Schematics of parallel bulk CRISPR screen and adapted *PerturbFate* Dual-omics Sequencing Library Preparation. **(A)** Schematics of *PerturbFate* bulk screen library preparation workflow. After genomic DNA extraction, the amplicon library was constructed for short reads sequencing. **(B)** Schematics of the *PerturbFate* Dual-omics workflow. After ligation, second-strand synthesis and read2 tagmentation on cDNA in-situ is performed. Referring to *ISSAAC-seq*(*21*), ATAC fragments are not affected by the second round of tagmentation due to the saturated Tn5 occupancy. Both ATAC and tagmented cDNA libraries are amplified in the multiplex PCR, while another round of sgRNA enrichment PCR is performed to fully barcode the sgRNA library. **(C)** Schematics of *PerturbFate* Dual-omics post-sequencing cell barcode matching. For each single cell with multiple molecular modalities, all modalities share the first two cell barcodes. The ATAC and steady-state transcriptome share the same cell barcode 3. For the sgRNA modality, each inner i7-barcode 4 combination corresponds to a unique barcode 3 pair from ATAC and RNA modalities. **(D)** Dot plot with kernel density estimation showing a high correlation in pseudobulk accessibility of shared peaks between single-modal *EasySci-ATAC*(*3*) and *PerturbFate* Dual-omics ATAC. HEK293T cells were profiled using both *EasySci-ATAC* and *PerturbFate* Dual-omics for benchmarking. Peaks were called separately and then merged for peak count quantification. Each library was normalized to 1e6. **(E)** Dot plot with kernel density showing a high correlation of gene expression profiles between single-modal *EasySci-RNA*(*3*) and *PerturbFate* Dual-omics RNA. HEK293T cells were profiled using both *EasySci-RNA* and *PerturbFate* Dual-omics for benchmarking. After aggregating single-cell gene counts, each library was normalized to 1e6. **(F)** Representative genome tracks demonstrating high consistency across modalities profiled using different techniques. RNA and ATAC tracks are shown at the same scale within their respective modalities

**Fig. S4:**
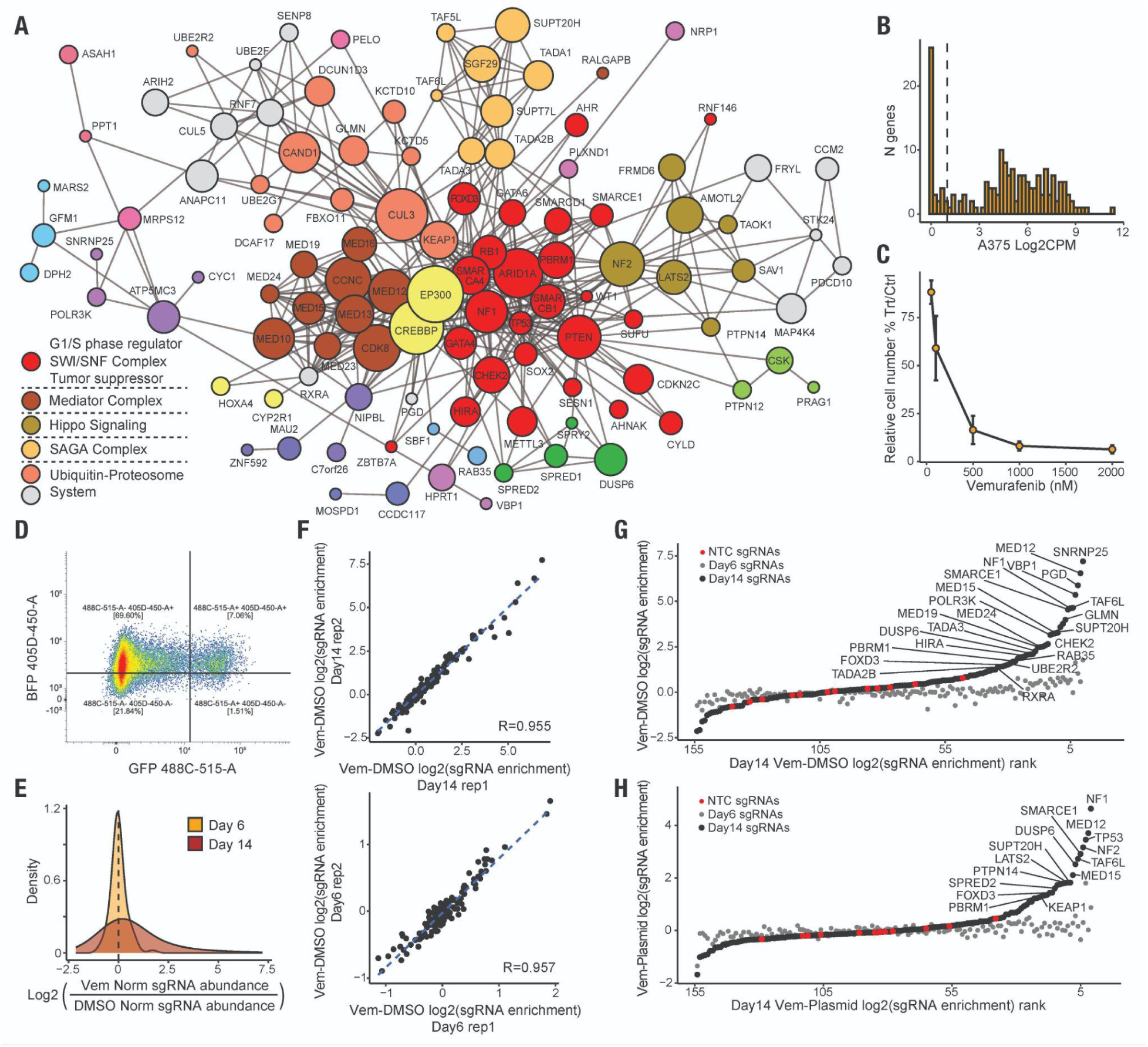
Selection and evaluation of the sgRNA library for *PerturbFate* screen. **(A)** Network graph of candidate Vemurafenib-resistant genes and main cluster annotations based on protein-protein interactions. Each dot represents a protein encoded by a candidate Vemurafenib-resistant gene, and each edge represents an interaction between two proteins. The size of the dots indicates the number of interactions detected within the network. Clustering was performed using MCL clustering in the STRING database(*54*), and the top functional annotations were extracted from the STRING database. **(B)** Histogram showing the expression distribution of candidate Vemurafenib-resistant genes in wild-type (WT) A375 cells. A375 cells were profiled using single cell RNA-seq, and gene counts across cells were aggregated. The pseudobulk gene counts were then normalized by the total gene count and multiplied by 1e6. A cutoff of Log2(CPM)=1 was used to filter out lowly expressed or non-expressed genes. **(C)** Dot plot with error bars showing melanoma cell growth inhibition by Vemurafenib treatment at different concentrations. A375 cells were seeded in a 6-well plate, and DMSO only (0 nM), 500 nM, 1000 nM, or 2000 nM Vemurafenib was added to the media of each well. After six days, the cell number in each well (n=2) was measured, and the relative cell number compared to cells grown in regular media was calculated. Error bars represent the mean ± standard deviation. **(D)** Representative flow cytometry gating strategy of selecting *dCas9-BFP-KRAB* expressing (BFP+) and sgRNA expressing (GFP+) cells. **(E)** Density plot showing the distribution of sgRNA abundance changes on Day 6 and Day 14 of the CRISPR screen. Normalized sgRNA abundance represents the relative abundance of each sgRNA pair compared to NTCs. The normalized sgRNA abundance ratio between Vemurafenib and DMSO conditions reflects the effect of genetic perturbations on cell growth under Vemurafenib treatment. **(F)** Dot plots showing the reproducibility of genetic perturbation effects across replicates of bulk CRISPR screen on day 14 (upper panel) and day 6 (lower panel). The Vem-DMSO log2(sgRNA enrichment) represents the normalized sgRNA abundance ratio between Vem and DMSO conditions, indicating the effect of genetic perturbations on cell growth under Vemurafenib treatment. **(G-H)** Dot plot showing the enrichment of top sgRNAs by comparing sgRNA abundance between Vem and DMSO conditions (**G**) or the plasmid library (**H**). The enrichment indicates whether a genetic perturbation enhanced cell growth under Vem treatment. Perturbations were ranked by their Day 14 enrichment, while Day 6 enrichment is shown in gray to indicate relatively unchanged sgRNA representation at this time point.

**Fig. S5:**
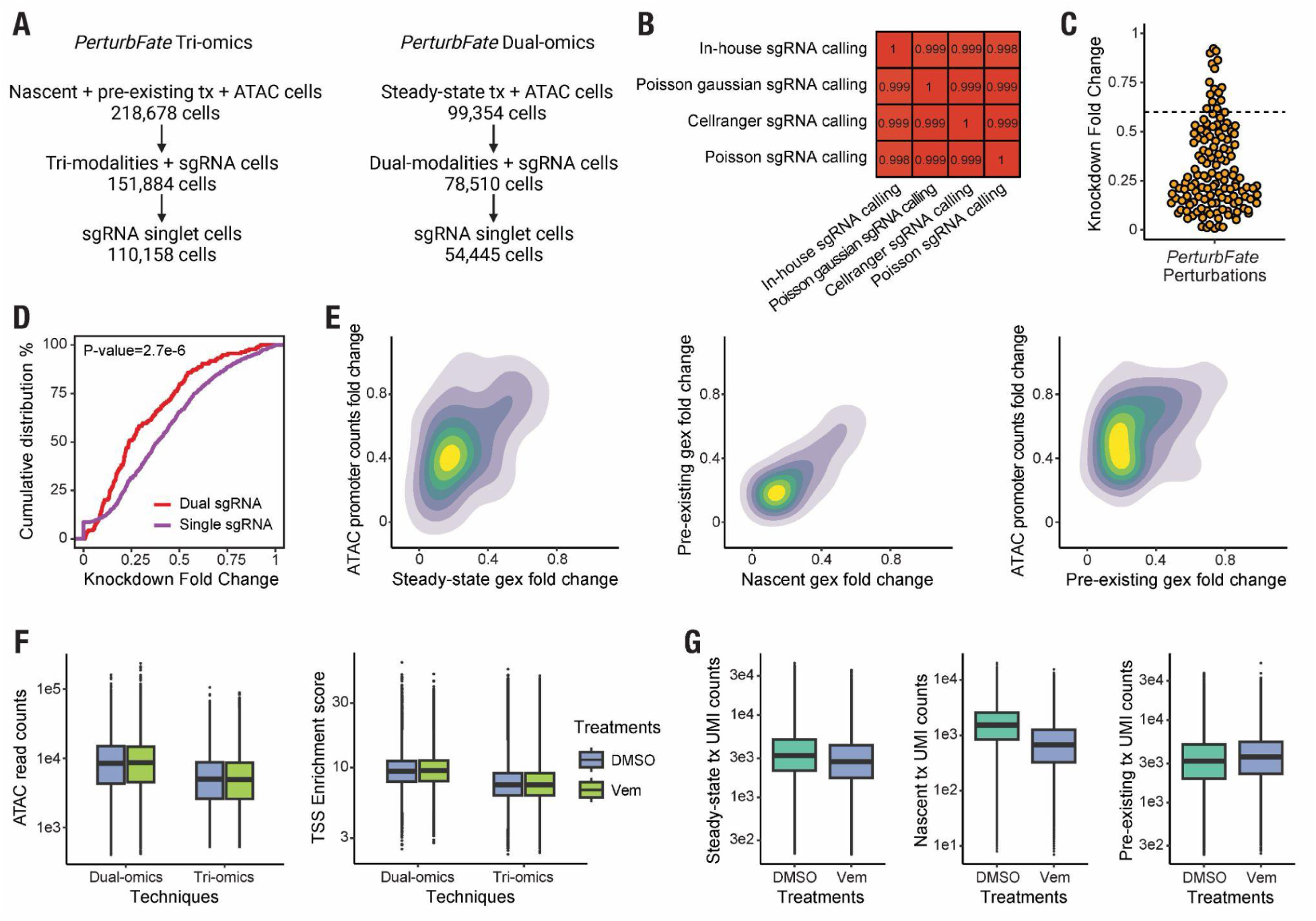
Evaluation and quality control of the *PerturbFate* library. **(A)** Schematics of computational processing of single cells generated from *PerturbFate* Tri-omics (Left) and *PerturbFate* Dual-omics (Right). **(B)** Heatmap comparing the cell number of sgRNA-assigned cell populations using different computational methods. Three metrics, total sgRNA counts, the proportion of primary sgRNA counts to total sgRNA counts, and the ratio of secondary to primary sgRNA counts, were used in the in-house sgRNA assignment method. Other statistical methods were implemented in crispat(*55*). **(C)** Beeswarm plot showing the fold change in steady-state expression of target genes compared to NTC. Gene counts from cells receiving the same sgRNA were aggregated and normalized to 1e6. The KD-to-NTC fold change of target genes was then calculated. **(D)** Cumulative distributions of knockdown gene expression fold change from *PerturbFate* using the dual-sgRNA system and *PerturbSci-Kinetics*(*8*) using the single-sgRNA system. A Kolmogorov-Smirnov (K-S) test was conducted to assess the statistical significance of the difference between distributions. Dual sgRNAs indeed exhibited a globally improved knockdown efficiency. **(E)** Density plots showing the consistency of target gene knockdown as reflected by different molecular modalities. To assess the knockdown effect at the ATAC layer, we identified all possible promoter regions of target genes spanning the TSS of all transcripts annotated in hg38. For each target gene, we retained only the promoter region that overlapped with the designed sgRNA target sites and exhibited the highest signal among all candidate promoter regions. The fold change in normalized ATAC counts within this region between KD cells and NTC cells was then used as a proxy for the loss of chromatin accessibility. **(F)** Box plots showing single-cell ATAC counts and TSS enrichment scores across techniques and conditions. TSS enrichment scores for single cells were calculated using SnapATAC2(*56*). **(G)** Box plots showing single-cell UMI counts for steady-state, nascent, and pre-existing transcriptomes across conditions. Boxes in box plots indicate the median and interquartile range (IQR), with whiskers indicating 1.5× IQR.

**Fig. S6:**
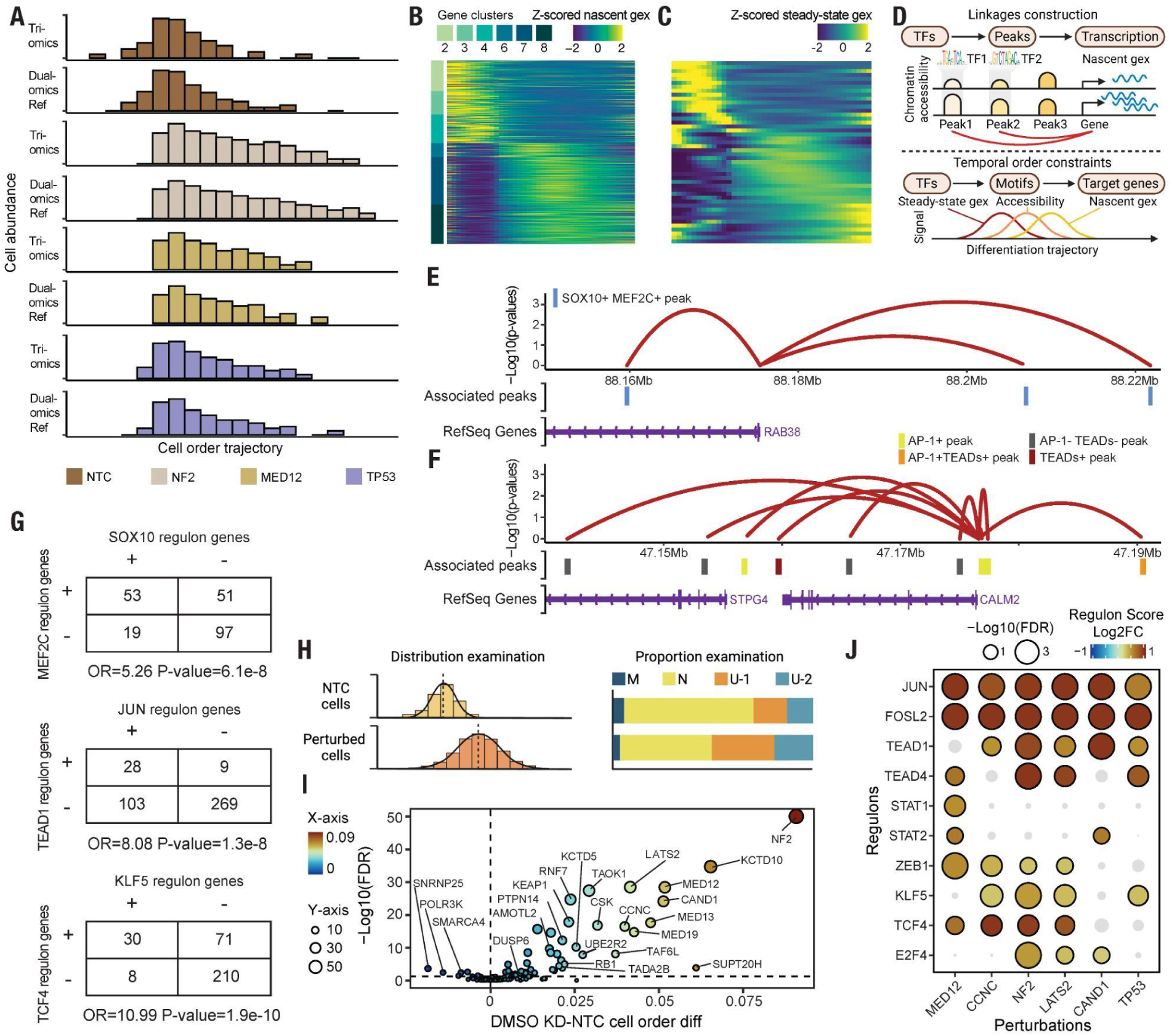
Genetic perturbations induced extensive cell state transitions through comprehensive remodeling of epigenetic landscape and gene expressions. **(A)** Histograms benchmarking the distributions of reference perturbed cell populations (Dual-omics) and projected cell populations (Tri-omics) from the DMSO condition along the cell order trajectory. To capture the transcriptomic phenotypes of perturbed A375 cells, the original diffusion map was generated using steady-state transcriptomes from perturbed single cells profiled by the Dual-omics protocol. To further infer cell state transition flows, single-cell nascent transcriptomes were incorporated by projecting similar pre-existing transcriptomes of Tri-omics cells onto the trajectory. Specifically, another diffusion map was generated by co-embedding single-cell whole transcriptomes and pre-existing transcriptomes. The nearest Tri-omics cell neighbor of each Dual-omics cell was then assigned to that cell in the original trajectory. The cell order of Tri-omics cells was inferred by averaging the cell orders of the three nearest Dual-omics cells. **(B)** Heatmap showing the dynamics of nascent gene expression along the trajectory. Genes are ordered identically to those in the steady-state gene expression heatmap in **Fig. 2F**. **(C)** Heatmap of steady-state gene expression profiles of TFs, ordered identically to those in **Fig. 2H**. These TFs exhibit significantly altered motif accessibility along the trajectory, with gene expression patterns that correspond to their motif accessibility changes. **(D)** Schematics of state transition temporal gene regulatory network reconstruction. TF-CRE-gene linkages were established by associating nascent gene expression with nearby ATAC peak counts and the presence of TF motifs within the associated peaks. Temporal constraints, based on the fundamental principles of transcriptional activation, were applied to refine linkages and select activating genes that follow the state transition direction. **(E)** A representative genome track showing CRE-gene linkages for *RAB38* and the embedding of SOX10 and MEF2C motifs within linked peaks. *RAB38* is significantly upregulated during the Neural crest-like to Melanocytic state transition, and the associated CREs of this gene contain both SOX10 and MEF2C motifs. **(F)** A representative genome track showing CRE-gene linkages for *CALM2* and the embedding of AP-1 and TEADs motifs within peaks. *CALM2* is significantly upregulated during the Neural crest-like to Undifferentiated-1 state transition, with AP-1 and TEAD motif occupancy identified in multiple associated ATAC peaks. **(G)** Example contingency tables showing significant co-regulation of TFs on state-specific genes. In the Melanocytic state, target genes identified by our strategy as being regulated by SOX10 and MEF2C exhibit significant overlap, consistent with a previous report(*43*). Similarly, JUN and TEAD1, as well as KLF5 and TCF4, showed strong overlaps in their target genes in the Undifferentiated-1 and Undifferentiated-2 states, respectively, aligning with previous studies(*47*, *57*). **(H)** Schematics of tests performed to quantify the effects of genetic perturbations on cell state transitions. K-S tests were conducted to compare the cell order distribution between KD cells and NTC cells, while proportion tests were used to assess the relative enrichment of KD cells in each cell state compared to NTC cells. **(I)** Volcano plot showing the difference in mean cell order between perturbed populations and NTC cells, along with the corresponding FDR-corrected K-S test significance. **(J)** Dot heatmap showing the increase in multi-modal regulon scores for representative perturbations and their statistical significance compared to NTC cells. To systematically assess regulon activation, we integrated nascent gene expression, associated peak accessibility, and TF steady-state gene expression into the multi-modal regulon score. Permutation testing was performed to compute the statistical significance of the change (**Methods**).

**Fig. S7:**
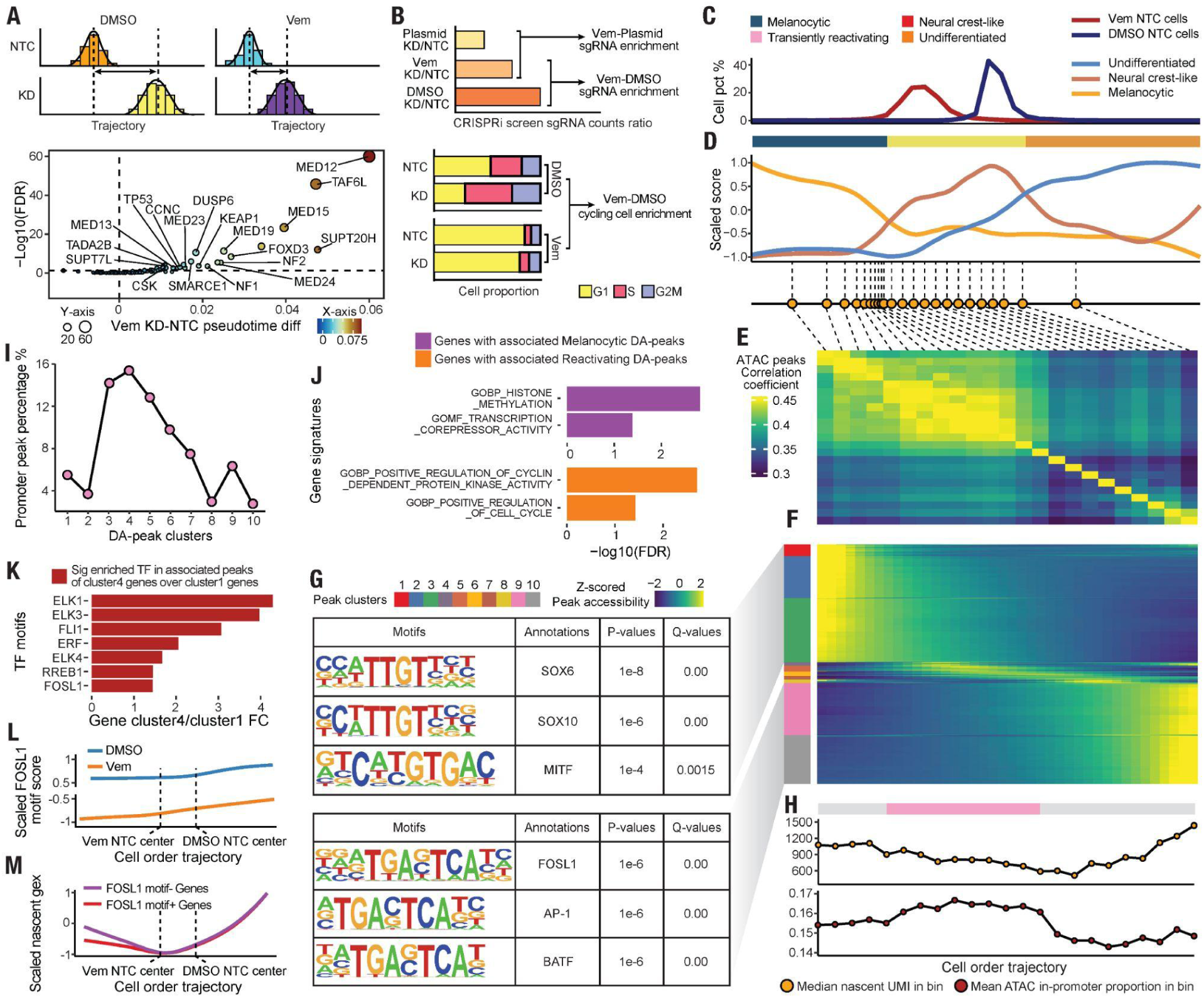
Comprehensive cell state transitions and epigenetic remodeling in response to Vemurafenib treatment. **(A)** Schematic comparing cellular transcriptomic state transitions between perturbed and NTC cells under two conditions (upper panel). The lower panel shows the volcano plot illustrating the difference in mean cell order between perturbed populations and NTC cells in the Vemurafenib treatment condition, along with the corresponding FDR-corrected p-values from K-S tests. **(B)** Schematic representations comparing sgRNA read counts between different conditions in parallel *PerturbFate* bulk CRISPR screens (upper panel) and comparing the proportion of cycling cells (inferred by steady-state transcriptome cell cycle score(*68*)) between perturbed cells and NTC cells across DMSO and Vemurafenib treatment conditions (lower panel). **(C)** Line plot showing the cell percentage distribution of NTC cells under DMSO and Vemurafenib treatment conditions along the cell order trajectory containing only Vemurafenib-treated cells. The full trajectory was inferred using cells from both DMSO and Vemurafenib-treated conditions, and Vemurafenib-treated cells were extracted for reconstructing this Vemurafenib treatment cell state transition trajectory. **(D)** Line plot showing melanoma phenotypic gene program scores along the cell order trajectory with only Vemurafenib-treated cells. The upper panel displays cell order cutoffs separating the Melanocytic, Neural crest-like and Undifferentiated states. The lower panel displays the averaged cell order within each bin throughout the trajectory. Given the strong effect of cell number on chromatin accessibility data sparsity, we prioritized an equal cell number per bin when binning the trajectory (**Methods**). **(E)** Heatmap showing the global chromatin accessibility landscape correlation between bins. Peak counts from cells within each bin were aggregated, normalized to 1e6, and used to calculate Pearson correlations. **(F)** Heatmap showing significant DA peaks at the bin pseudobulk level. Binned pseudobulk ATAC peak counts and the average cell order of bins were used to identify DA peaks. DA peaks from clusters 1-3 were defined as Vem-sensitive Melanocytic state DA peaks, while DA peaks from clusters 9-10 were defined as Vem-resistant Undifferentiated state DA peaks. Motif enrichment was performed using HOMER(*69*). **(G)** Tables showing the top enriched TF motifs in DA peaks opening in the Vem-sensitive Melanocytic state and Vem-resistant Undifferentiated state. DA peaks from clusters 1-3 were defined as Vem-sensitive Melanocytic state DA peaks, while DA peaks from clusters 9-10 were defined as Vem-resistant Undifferentiated state DA peaks. Motif enrichment was performed using HOMER(*69*). **(H)** Line plots with dots showing median nascent UMI counts per cell within each bin (upper panel) and mean ATAC in-promoter read proportion per cell within each bin (lower panel) along the trajectory. In the Vemurafenib-inhibited state, we observed the most suppressed transcriptional and promoter activities. As cells were transiting from this state to the Vem-sensitive Melanocytic state, a sharp wave of promoter region opening was detected, consistent with the global activation of motif accessibility (**Fig. 3J**), and we defined this state as the Transiently reactivating state. The upper annotation bar indicates cell order cutoffs for defining the Transiently reactivating state; Grey areas represent undetermined states in this panel. **(I)** Line plot with dots showing the proportion of DA peaks within promoter regions in each cluster along the trajectory. The increase in in-promoter DA peaks during the reactivation stage aligns with observations in **Fig. S7D-J,** Bar plots showing top functional enrichments for genes with associated peaks in clusters 1-3 DA peaks (Vem-sensitive Melanocytic state DA peaks) or clusters 4-6 DA peaks (Vem-sensitive Reactivating state DA peaks). The enrichment of histone modification-related terms in genes associated with Vem-sensitive Melanocytic state DA peaks further supports the involvement of epigenetic remodeling in melanoma cell state transitions. The enrichment of cell cycle machinery-related terms in genes associated with Vem-sensitive reactivating DA peaks corroborates gene regulation reactivation following the Vemurafenib-inhibited state. **(K)** Bar plots showing the enrichment fold of TF motif occurrences in associated peaks between genes that show maximum upregulation in the Vem-resistant Undifferentiated state (cluster 4 genes) and genes that show maximum upregulation in the Vem-responsive Reactivating state (cluster 1 genes) (**Fig. 3I**). After identifying the embedding of significant DA motifs (**Fig. 3J**) in genes, proportion tests were performed. **(L)** Line plot showing the scaled motif chromVAR score of FOSL1 in cells from the DMSO and Vemurafenib treatment conditions along the integrative trajectory. FOSL1 motif accessibility exhibited a consistent increase toward the Undifferentiated state. **(M)** Line plot showing the averaged scaled nascent expression trend of Vem-resistant GRN member genes with or without FOSL1 motifs in their associated peaks. The lack of activity of FOSL1 target genes in the Melanocytic state supports its essential role in driving melanoma dedifferentiation and drug resistance.

**Fig. S8:**
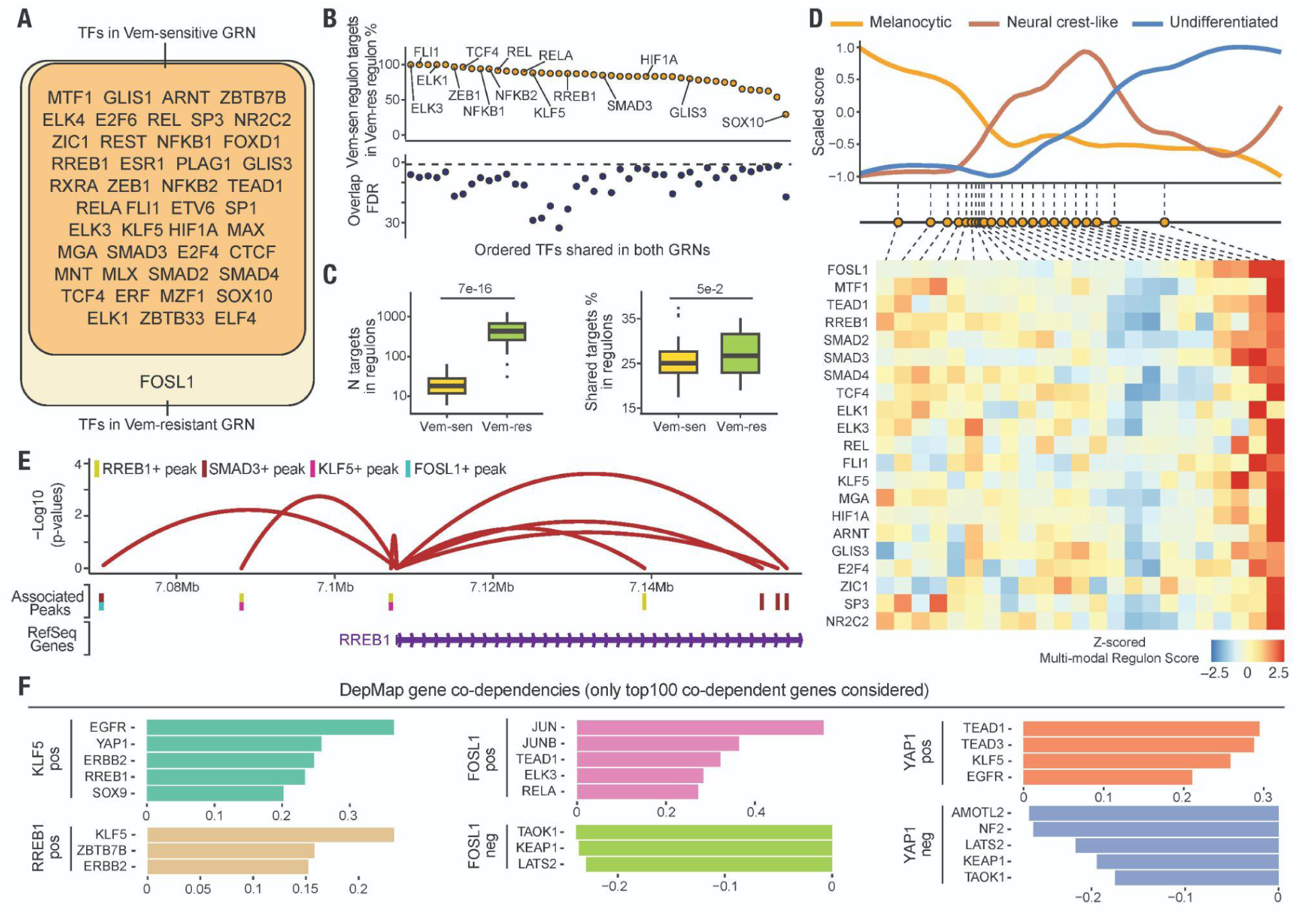
The establishment of state-transiting gene regulatory networks under the Vemurafenib treatment. **(A)** Venn diagram showing the overlap of TF regulators in the Vem-sensitive and Vem-resistant GRNs. Only FOSL1 is within the motif cluster exhibiting a single peak at the Vem-resistant Undifferentiated state. **(B)** Dot plots showing the percentage of Vem-sensitive GRN target genes that are also target genes of the same TF regulator in the Vem-resistant GRN (upper panel). The lower panel displays the FDR-corrected statistical significance of the overlap calculated by Fisher’s exact tests. **(C)** Box plots comparing the number of target genes per regulon (left panel) and the average percentage of shared target genes among regulons (right panel) between Vem-resistant and Vem-sensitive GRNs. The presence of two waves of gene regulation activation (**Fig. 3I-J**), along with their strong similarities in gene expression and epigenetic landscapes (**Fig. 3I-J**, **Fig. S8B**), the significantly smaller number of target genes per regulon, and the significantly lower collaboration between regulons in the Vem-sensitive GRN, and the strong overlap of Vem-sensitive regulon target genes within Vem-resistant regulons, all support the notion that the Vem-sensitive GRN represents a partially activated sub-network of the Vem-resistant GRN. **(D)** Heatmap showing z-scored multi-modal regulon scores within each bin along the trajectory. A unified pattern of the strongest inhibition at the Vem NTC center and the strongest activation at the Vem-resistant Undifferentiated state was observed. **(E)** A representative genome track showing CRE-RREB gene linkages and the embedding of RREB1, FOSL1, SMAD3, and KLF5 motifs within peaks. **(F)** Bar plots showing co-dependencies between key genes involved in melanoma drug resistance identified by DepMap(*66*). Only the top 100 co-dependent genes of the query gene were considered. Boxes in box plots indicate the median and interquartile range (IQR), with whiskers indicating 1.5× IQR.

**Fig. S9:**
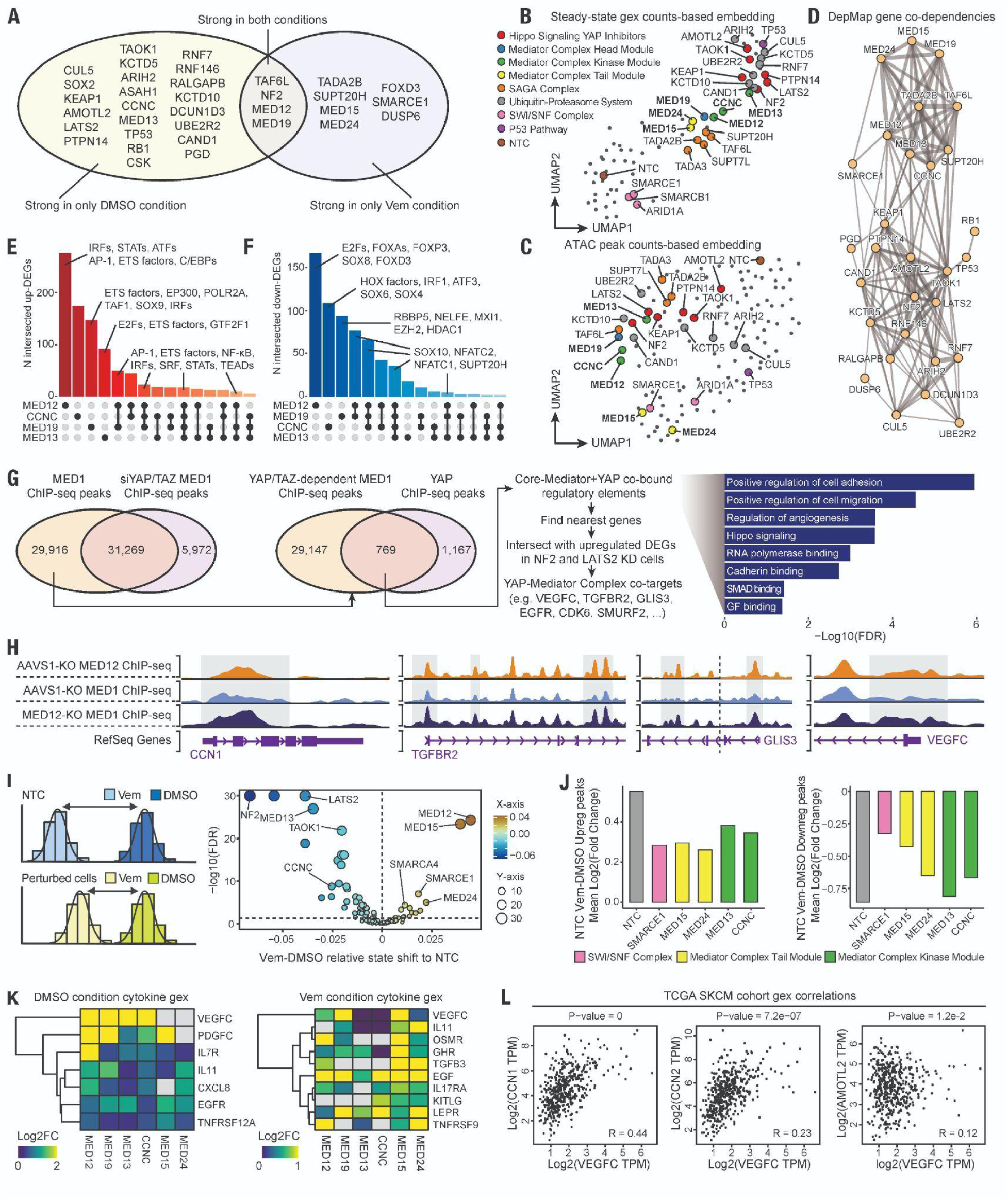
Perturbation of Mediator Complex genes in melanoma induces heterogeneous phenotypic responses, with involvement of a converged Hippo/YAP-Mediated gene program. **(A)** Venn diagram showing the overlap of genetic perturbations that exhibit strong dedifferentiation state shifts relative to NTC between DMSO and Vemurafenib treatment conditions. **(B-C)** UMAP embeddings of pseudobulk steady-state transcriptomes (**B**) and pseudobulk open chromatin landscapes (**C**) of genetic perturbations in the DMSO condition, highlighting structure-concordant Mediator Complex sub-module functionalities. Perturbed genes are colored by their primary function. While Mediator Complex perturbations cluster closely, confirming their shared role as a complex, Mediator Complex Kinase Module-associated perturbations are positioned near the Hippo signaling YAP inhibitor perturbation cluster, indicating shared perturbation response features. **(D)** Network graph showing co-dependencies between strong perturbations identified through the DepMap pan-cancer bulk CRISPR screen. Only the top 100 co-dependent genes of the query gene are included. The width of edges represents the correlation between genes. The co-dependencies between Mediator Complex Kinase Module genes and Hippo signaling YAP inhibitor genes further validated their potential converged functionalities in cancer. **(E-F)** Bar plots illustrating the overlap of upregulated (**e**) or downregulated (**F**) steady-state transcriptome DEGs between Mediator Complex Kinase Module perturbations, and their TF motif enrichments within promoter regions identified by RcisTarget. Shared upregulated DEGs between *MED12* and *MED19* perturbations exhibit strong promoter enrichments in AP-1 and NFKB, while the strongest SOX10 enrichment is observed in shared downregulated DEGs between *MED12* and *MED19*. **(G)** Workflow for identifying YAP-Mediator Complex co-regulatory genomic regions and target genes. Original ChIP-seq data were obtained from(*74*), and GREAT(*81*) was used to identify the nearest genes and annotate functional enrichments of regulatory regions. **(H)** Representative genome tracks showing co-occupancy patterns between the Core-Mediator (represented by MED1) and the Mediator Kinase Module, as well as the derepression of Core-Mediator binding upon Mediator Kinase Module knockout on YAP-Mediator Complex co-target genes. Representative binding peaks are highlighted by grey rectangles. Original ChIP-seq data were obtained from(*72*). **(I)** Schematics compare the distance of cell order transition upon Vemurafenib treatment between NTC and perturbed cells (**Methods**). The right panel shows the relative resistance or hypersensitivity of perturbations to Vemurafenib treatment compared to NTC and corresponding statistical significance (FDR-corrected two-sided p-values from Wilcox tests). Except for *MED12*-perturbed cells, knocking down *MED15* and *MED24*, and the chromatin remodeler *SMARCE1* showed the highest extent of resistance. **(J)** Bar plots show the averaged log2 fold change in ATAC counts of NTC-upregulated (left panel) and NTC-downregulated (right panel) peaks across different perturbations upon Vemurafenib treatment. Compared to *CCNC*- and *MED13*-perturbed cells, *MED15*- and *MED24*-perturbed cells exhibited a less altered epigenetic landscape after Vemurafenib treatment. **(K)** Heatmaps showing steady-state cytokine gene expression in each treatment condition. To identify potential cytokines involved in Mediator Complex perturbation-induced phenotypic state transitions, only cytokine genes with significant activation in Mediator Complex Kinase Module perturbations in the DMSO condition and cytokine genes with significant activation in Mediator Complex Tail Module perturbations in the Vemurafenib treatment condition were included. Among them, *VEGFC* is the only cytokine gene that consistently exhibits activation patterns aligning with the observed phenotypic state transition across conditions. **(L)** Dot plots showing the correlation between *VEGFC* expression and representative Hippo/YAP signaling target genes (*i.e., CCN1, CCN2, AMOTL2*) in the TCGA SKCM cohort.

**Fig. S10:**
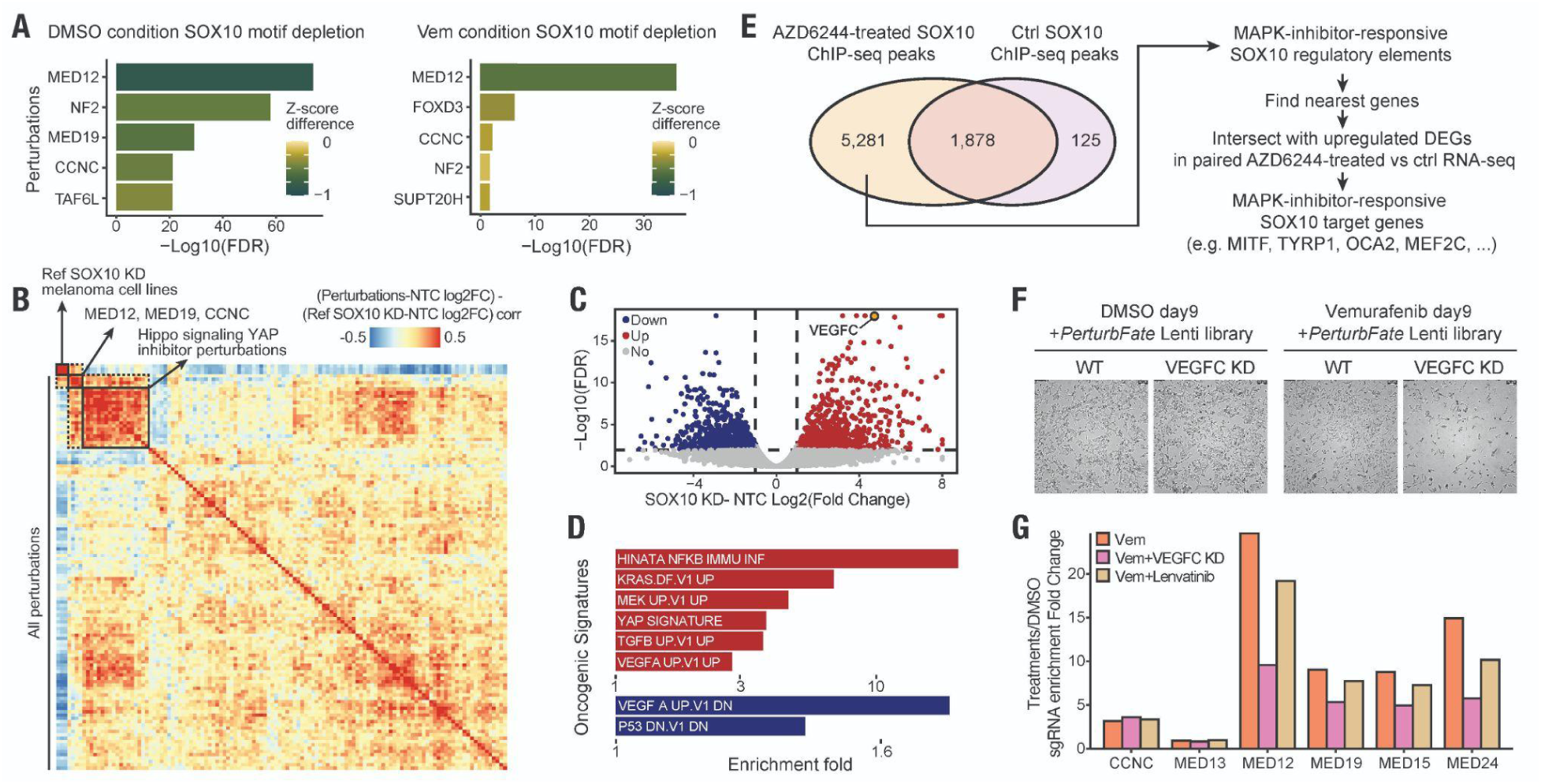
*MED12* knockdown partially mimics the SOX10 loss-of-function, and the validation of *VEGFC* to be a strong convergent mediator of Mediator Complex perturbations induced-Vemurafenib resistance. **(A)** Bar plots showing the top perturbations with the strongest depletion of SOX10 motifs and their statistical significance. Mediator Complex Kinase Module perturbations, particularly *MED12*, resulted in the most significant depletion of SOX10 accessibility. Additionally, the persistent depletion of SOX10 activity after Vemurafenib-induced melanoma differentiation in *MED12* KD strongly suggests impaired SOX10 function. **(B)** Heatmap showing the correlation of gene expression changes induced by *SOX10* KD in melanoma cell lines (originally profiled in(*75*)) and all candidate drug-resistant gene KDs examined in our study (**Methods**). Hierarchical clustering identified high intra-category similarities among Mediator Complex Kinase Module and Hippo signaling YAP inhibitor perturbations. While most perturbations induced transcriptomic changes distinct from *SOX10* KD, and perturbations driving melanoma dedifferentiation (e.g., Hippo signaling YAP inhibitor perturbations) didn’t cause complete SOX10 loss-of-function, Mediator Complex Kinase Module perturbations closely mimic the *SOX10* KD phenotype. **(C)** Volcano plot showing gene expression changes upon *SOX10* KD in multiple melanoma culture models. DEGs were identified based on an FDR-corrected two-sided p-value < 0.01 and an absolute fold change ≥ 1. To account for melanoma culture model-specific effects, cell line identities were included as covariates in the DEG analysis. *VEGFC* was identified as a top shared upregulated gene upon *SOX10* KD across melanoma models. **(D)** Functional enrichment of oncogenic signatures in upregulated and downregulated genes upon *SOX10* KD. The recurrent enrichment of MAPK, YAP, and VEGF signatures further confirmed the convergence of downstream gene programs between *SOX10* KD and Mediator Complex Kinase Module perturbations in melanoma. **(E)** Workflow illustrating the identification of genomics regions bound by SOX10 and SOX10 target genes that are induced by another MAPKi (AZD6244) treatment in melanoma. In addition to a subset of housekeeping SOX10 binding sites in the chromatin, MAPKi treatment strongly induced SOX10 activity. By intersecting the nearest genes of SOX10 peaks with upregulated DEGs in paired RNA-seq data, we identified direct SOX10 target genes induced upon MAPKi treatment, including key melanogenesis genes. **(F)** Microscopic images showing cell growth of *VEGFC* KD or WT cell pools with *PerturbFate* perturbations to develop Vemurafenib resistance. VEGFC loss-of-function resulted in a strong global reduction in cell growth under Vemurafenib treatment but had no observable effects in the DMSO condition. An equal number of cells was seeded in each plate on day 0, and cells in the DMSO condition were 1:3 split once during the culture. **(G)** Bar plot showing the relative growth advantage of Mediator Complex-perturbed cells in the Vemurafenib treatment condition compared to the DMSO condition, as well as their reduced drug resistance upon *VEGFC* KD or Lenvatinib treatment, a VEGFR inhibitor. The fold change of the Vem+*VEGFC* KD group was averaged from three replicates. Consistent with our analyses, CCNC and MED13 perturbations resulted in minimal or no acquired drug resistance, and their growth was unaffected by either genetic or chemical inhibition of the VEGFC pathway. Although Lenvatinib treatment had mild effects, potentially due to suboptimal dosing or the timing of inhibition, its effect was consistent in direction with *VEGFC* KD.

