## Supplementary Text 1 for "Uncovering Convergent Cell State Dynamics Across Divergent Genetic Perturbations Through Single-Cell High-Content CRISPR Screening"

***PerturbFate* costs estimation**

| Steps | Purposes | Reagents | Unit cost (USD/unit) | Overall cost in experiment (USD) | |
| --- | --- | --- | --- | --- | --- |
| Upstream processing steps | | | | | |
| Fixation | PBSI | SUPERase In | 8.43/1M cells | 16.86 | 18.5 |
|  |  | 25x Protease Inhibitor | 0.1255/1M cells | 0.251 |  |
|  |  | BSA (NEB) | 0.6375/1M cells | 1.275 |  |
|  | Cell fixation | 16% Formaldehyde | 0.054/1M cells | 0.108 |  |
|  |  | 10% Triton-X100 | 0.0014/1M cells | 0.0028 |  |
| ATAC in bulk | 4xTD buffer | DMF | 0.0062/50k cells | 0.074 | 26.63 |
|  |  | KAc | 0.00081/50k cells | 0.0097 |  |
|  |  | MgAc | 0.00067/50k cells | 0.008 |  |
|  | Other reagents in the reaction | SUPERase In | 0.562/50k cells | 6.744 |  |
|  |  | RNaseOUT | 0.948/50k cells | 11.376 |  |
|  |  | 10% Triton-X100 | 0.000014/50k cells | 0.000168 |  |
|  |  | 25x Protease Inhibitor | 0.0083/50k cells | 0.1 |  |
|  | PerturbFate-ATAC Tn5 | Tn5 | 0.693/50k cells | 8.32 |  |
| Cell washing | PBB-0.05 | BSA (NEB) | 0.85/mL buffer | 8.5 | 23.31 |
|  |  | 10% Triton-X100 | 0.00028/mL buffer | 0.0028 |  |
|  | PBB-0.1 | BSA (NEB) | 0.85/mL buffer | 2.55 |  |
|  |  | 10% Triton-X100 | 0.00056/mL buffer | 0.0017 |  |
|  | PSBI | SUPERase In | 11.24/mL buffer | 11.24 |  |
|  |  | BSA (NEB) | 0.85/mL buffer | 0.85 |  |
|  |  | 25x Protease Inhibitor | 0.1672/mL buffer | 0.1672 |  |
| Reverse Transcription in bulk | Primers master mix | dNTP (10mM each) | 1.932/96well plate | 7.728 | 330.37 |
|  |  | dT primers (100uM) | 1.261/96well plate | 5.043 |  |
|  |  | rN primers (100uM) | 1.229/96well plate | 4.916 |  |
|  |  | sgRNA primer (10uM) | 0.227/96well plate | 0.909 |  |
|  | RT reaction master mix | 5xPEG NaCl RT buffer | 0.614/96well plate | 2.456 |  |
|  |  | Maxima H Minus RTase | 61.875/96well plate | 247.5 |  |
|  |  | SUPERase In | 15.455/96well plate | 61.82 |  |
| Cell washing | Reaction quenching | EDTA (500mM) | 0.024/96well plate | 0.0962 | 0.0962 |
| The 1^st^ ligation | Ligation barcoding | The 1^st^ ligation adapters | 0.919/96well plate | 3.676 | 275.684 |
|  | T4 ligase reaction mix | T4 Ligase | 61.82/96well plate | 247.28 |  |
|  |  | SUPERase In | 6.182/96well plate | 24.728 |  |
| The 2^nd^ ligation | Ligation barcoding | The 2^nd^ ligation adapters | 0.678/96well plate | 2.713 | 274.721 |
|  | T4 ligase reaction mix | T4 Ligase | 61.82/96well plate | 247.28 |  |
|  |  | SUPERase In | 6.182/96well plate | 24.728 |  |
| In-situ Click Chemistry | Click Chemistry Reaction | Copper protectant | 12.52/rxn | 50.08 | 72.99 |
|  |  | Biotin-picolyl-azide (10mM) | 0.108/rxn | 0.431 |  |
|  |  | SUPERase In | 5.62/rxn | 22.48 |  |
| Total cost of PerturbFate upstream protocol: 1022.3 USD  Cells processed in total: 384,000 cells  Cost per cell: 0.00266/cell | | | | | |
| Tri-omics downstream processing steps | | | | | |
| Cell lysis | Cell lysis buffer | Proteinase K (20mg/ml) | 0.1135/PCR strip | 1.816 | 1.8541 |
|  |  | 1% SDS | 0.00074/PCR strip | 0.01184 |  |
|  | Stop buffer | 10% Tween-20 | 0.000784/PCR strip | 0.012544 |  |
|  |  | PMSF (100mM) | 0.000858/PCR strip | 0.013728 |  |
| Strep-biotin pulldown | Pulldown reaction | MyOne C1 strep beads | 11.04/PCR strip | 176.64 | 375.33 |
|  |  | 1xBW-T buffer | 0.021/PCR strip | 0.3371 |  |
|  |  | 2xBW buffer | 0.006125/PCR strip | 0.098 |  |
|  |  | SUPERase In | 6.6316/PCR strip | 106.106 |  |
|  | Beads washing | 1xBW-T buffer | 0.0268/PCR strip | 0.4291 |  |
|  |  | SUPERase In | 5.7325/PCR strip | 91.72 |  |
| Second Strand Synthesis on beads | SSS master mix | SSS buffer | 18.975/PCR strip | 303.6 | 770.7 |
|  |  | SSS enzyme |  |  |  |
| Second Strand Synthesis in supernatant | Purification on dsDNA and cDNA-RNA hybrid | RNAclean XP beads | 10.22/PCR strip | 163.5 |  |
|  | SSS master mix | SSS buffer | 18.975/PCR strip | 303.6 |  |
|  |  | SSS enzyme |  |  |  |
| Read2 tagmentation | Read2 tagmentation on nascent tx | Nextera Read2 Tn5 | 0.035/PCR strip | 0.56 | 12.546 |
|  | Read2 tagmentation on pre-existing tx | Nextera Read2 Tn5 | 0.069/PCR strip | 1.104 |  |
|  | Rxn inactivation | BSA (NEB) | 0.34/PCR strip | 10.88 |  |
|  |  | 1% SDS | 0.00006/PCR strip | 0.0019 |  |
| Multiplex PCR | PCR rxn | Universal P5 primer | 0.1571/PCR strip | 5.0272 | 425.203 |
|  |  | Inner i7 U6 primer | 0.0896/PCR strip | 2.8672 |  |
|  |  | Indexed Nextera R2-P7 primer | 0.0809/PCR strip | 2.5888 |  |
|  |  | NEBnext 2x PCR MM | 12.96/PCR strip | 414.72 |  |
| sgRNA enrichment PCR | PCR rxn | Universal P5 primer | 0.0197/PCR well | 0.3152 | 26.397 |
|  |  | Indexed TruSeq R2-P7 primer | 0.0101/PCR well | 0.1616 |  |
|  |  | NEBnext 2x PCR MM | 1.62/PCR well | 25.92 |  |
| DNA purification | sgRNA library column purification | DNA clean&concentrator-5 | 1.716/column | 1.716 | 65.646 |
|  | Main libraries column purification | DNA clean&concentrator-5 |  | 3.431 |  |
|  | sgRNA library gel extraction | Zymoclean gel DNA recovery | 1.93/column | 1.93 |  |
|  | Main libraries gel extraction | Zymoclean gel DNA recovery |  | 3.86 |  |
| Total cost of Tri-omics downstream protocol: 1623 USD  Cells processed by Tri-omics downstream protocol: 256,000 cells  Cost per cell: 0.00634 USD/cell | | | | | |
| Dual-omics downstream processing steps | | | | | |
| Second Strand Synthesis | SSS master mix | SSS buffer | 9.4875/PCR strip | 75.9 | 75.9 |
|  |  | SSS enzyme |  |  |  |
| Read2 tagmentation | Read2 tagmentation on steady-state tx | Nextera Read2 Tn5 | 0.069/PCR strip | 0.552 | 2.197 |
|  | Rxn inactivation and uncrosslinking | Proteinase K (20mg/ml) | 0.2043/PCR strip | 1.6344 |  |
|  |  | 1% SDS | 0.001332/PCR strip | 0.0107 |  |
| Multiplex PCR | PCR rxn | Universal P5 primer | 0.1571/PCR strip | 1.2568 | 106.301 |
|  |  | Inner i7 U6 primer | 0.0896/PCR strip | 0.7168 |  |
|  |  | Indexed Nextera R2-P7 primer | 0.0809/PCR strip | 0.6472 |  |
|  |  | NEBnext 2x PCR MM | 12.96/PCR strip | 103.68 |  |
| sgRNA enrichment PCR | PCR rxn | Universal P5 primer | 0.0197/PCR well | 0.1576 | 13.20 |
|  |  | Indexed TruSeq R2-P7 primer | 0.0101/PCR well | 0.0808 |  |
|  |  | NEBnext 2x PCR MM | 1.62/PCR well | 12.96 |  |
| DNA purification | sgRNA library column purification | DNA clean&concentrator-5 | 1.716/column | 1.716 | 7.292 |
|  | Main libraries column purification | DNA clean&concentrator-5 |  | 1.716 |  |
|  | sgRNA library gel extraction | Zymoclean gel DNA recovery | 1.93/column | 1.93 |  |
|  | Main libraries gel extraction | Zymoclean gel DNA recovery |  | 1.93 |  |
| Total cost of Dual-omics downstream protocol: 204.89 USD  Cells processed by Tri-omics downstream protocol: 128,000 cells  Cost per cell: 0.0016 USD/cell | | | | | |

Final cost for each cell of PerturbFate (Tri-omics): 0.00266+0.00634=**0.009 USD**

Final cost for each cell of PerturbFate (Dual-omics): 0.00266+0.0016=**0.00426 USD**
